# Characterization of aperiodic and theta activity in preterm infants using EEG: Insights into cerebral maturation and inter-individual variability

**DOI:** 10.64898/2025.12.26.696574

**Authors:** Aline Gonzalez-Carpinteiro, Hala Nasser, Amandine Pedoux, Laurie Devisscher, Kunyan Cai, Nicolas Elbaz, Chloé Ghozland, Léa Guéret, Sara Neumane, Yann Leprince, Aline Lefebvre, Lucie Hertz-Pannier, Alice Heneau, Alice Frérot, Olivier Sibony, Catherine Delanoë, Marianne Barbu-Roth, Marianne Alison, Richard Delorme, Valérie Biran, Jessica Dubois, Parvaneh Adibpour

## Abstract

Disrupting critical processes of brain development, preterm birth interferes with the maturation of brain networks and functional activity, including theta oscillations that are thought to play a key role in early network formation. Traditional EEG spectral analyses have indicated marked development of theta power in early infancy, but these approaches mix oscillatory and non-oscillatory activity, limiting insights into the mechanisms underlying the neural changes. Using spectral parameterization, we aimed to evaluate developmental changes in aperiodic activity and periodic theta power in infants born very preterm compared to full-terms, and to further explore whether clinical factors and brain microstructure could explain the inter-individual variability within preterms.

High-density EEG was acquired during active/REM sleep at term-equivalent age (TEA) and 2 months corrected age (2mCA) in 41 very preterm infants (born <32 weeks gestational age [GA]; mean ± standard deviation: 26.9±1.7 weeks) and 13 full-term controls (born ≥37 weeks GA; 40.1±1 weeks). Spectral parameterization was used to extract aperiodic components (offset, exponent) and periodic theta power globally and across spatial clusters of electrodes (anterior, central, posterior). From TEA to 2mCA, offset, exponent, and theta power increased with no differences between preterm and full-term infants. At TEA, metrics of aperiodic activity were stronger in anterior compared with posterior areas, but this regional landscape shifted by 2mCA with pronounced increases in aperiodic offset and exponent in posterior areas from TEA to 2mCA. Within preterms, inter-individual variability in aperiodic and periodic activity at TEA was partly explained by clinical risk factors: male sex, lower gestational-age at birth, small weight at birth, and invasive ventilation were linked to alterations of aperiodic offset, exponent and theta power. Additionally, higher theta power at TEA correlated with lower cortical fractional anisotropy assessed with diffusion MRI at the same age, consistently with more advanced maturation of the brain. Collectively, these findings indicate that EEG spectral parameterization combined with spatial analysis provides a sensitive framework for characterizing the postnatal maturation of brain activity in infants, as well as early vulnerabilities associated with prematurity and perinatal adversity.

## Introduction

Preterm birth affects 13.4 million infants (∼10% of live birth) each year (Ohuma et al., 2023) and is associated with long-term neurodevelopmental atypicalities (Bradley et al., 2025; World Health Organization, 2023). It occurs during a period (the last trimester of pregnancy) when brain development follows a highly orchestrated sequence of neurobiological processes, including neuronal migration, synaptogenesis, growth of connections and establishment of cerebral networks (Kostović & Judaš, 2015). Premature birth interrupts these processes and increases vulnerability to atypical neurodevelopmental trajectories, with consequent long term difficulties in cognitive, sensory and motor domains (Ancel et al., 2006; Johnson & Marlow, 2017; Ream & Lehwald, 2018), particularly when the birth is very premature, i.e. before 32 weeks of gestational age (GA) (Pierrat et al., 2021). Studying the early trajectories of brain development in preterm infants is therefore essential for understanding prematurity-related alterations in cerebral functioning and for identifying early markers of atypical outcome.

Clinical evaluations of neonatal brain development often involve non-invasive electroencephalography (EEG) recordings obtained during sleep. These recordings allow clinicians to qualitatively assess how brain activity is organized across brain regions and over time throughout sleep cycles, helping to detect markers of brain maturation or abnormal functioning (Wallois et al., 2021; Bourel-Ponchel et al., 2021; Iyer et al., 2015; André et al., 2010; Vanhatalo & Kaila, 2006; Mirmiran et al., 2003). Such assessments have further helped to identify the neonatal precursors of rapid eye movement (REM) sleep (∼active sleep) and non-REM sleep, which both exhibit characteristic oscillatory activity in particular within the delta (1-3 Hz) and theta (4-7 Hz) frequency bands (Dereymaeker et al., 2017; Grigg-Damberger, 2016). Animal models have suggested that these oscillatory activity patterns, and in particular theta rhythm during REM sleep, play an important role in the development of functional networks. Specifically, they are thought to promote long-range connectivity, including connectivity between cortical and sub-cortical structures, contribute to plasticity mechanisms, and coordinate activity within developing sensorimotor circuits, thereby supporting the early integration of sensorimotor networks during REM sleep (Brockmann et al., 2011; Del Rio-Bermudez et al., 2017; Del Rio-Bermudez & Blumberg, 2018).

In human infants, several EEG studies have quantitatively investigated early sleep activity through spectral analyses by extracting the power of the recorded signals within different frequency bands (e.g. Novelli et al., 2016; Sankupellay et al., 2011). In particular, an increase in theta band power during active/REM sleep has been described from the late preterm period (i.e., ∼34 weeks GA) up to approximately 1 month after term-equivalent age (TEA) (Corsi-Cabrera et al., 2020; Whitehead et al., 2018). This developmental increase in theta power appears to parallel the transition from early diffuse bursting activity towards more refined cortical functional dynamics (Yrjölä et al., 2024). Study of theta activity can be a particularly informative framework for understanding vulnerable populations such as preterm infants, in whom sensorimotor networks are known to be especially affected by early adversity (Devisscher, 2026; Neumane, 2022), and in light of the beforementioned relevance of theta activity for the development of sensorimotor networks. Consistent with this, in very preterm infants, an altered theta-band power and connectivity has been reported at TEA when compared to full-term neonates (Guyer et al., 2019; Tokariev et al., 2016; Yrjölä et al., 2022), with some alterations persisting up to 3 months of corrected age (mCA) (Guyer et al., 2019). At this age, preterms display more pronounced occipital than central distribution of theta activity, indicating that prematurity exerts heterogeneous, region-specific effects on early brain activity (Guyer et al., 2019)

However, traditional spectral power measures (e.g., absolute power) mix oscillatory and non-oscillatory background neural activity (Donoghue, Dominguez, et al., 2020). Recent spectral parameterization techniques address this limitation by decomposing the power spectrum into two components: an aperiodic component, reflecting the non-oscillatory background neural activity that follows a 1/f-like pattern, and periodic components reflecting rhythmic oscillations manifested as peaks rising above the 1/f distribution. The aperiodic component is described by an offset, representing baseline power level, and an exponent, indicating the slope of power decay across frequencies (Donoghue, Dominguez, et al., 2020; Donoghue, Haller, et al., 2020). Absolute power reflects the contribution of both components, and a change in band power can result from a true change in oscillatory activity (e.g., a stronger or more frequent rhythm), a change in the underlying aperiodic background (e.g. a shift in the overall power level or in the slope of spectral decay), or a combination of both (Donoghue, Haller, et al., 2020). Yet distinguishing aperiodic and oscillatory activity is particularly important in developmental research, as underlying developmental mechanisms may differ (Donoghue et al., 2020; Schaworonkow & Voytek, 2021). In addition, oscillatory frequencies can vary across individuals and developmental stages (Schaworonkow & Voytek, 2021; Wilkinson et al., 2024). For example, a developmental increase in theta band power (quantified with traditional spectral analysis) could reflect a genuine rise in oscillatory theta activity (indexing the maturation of long-range network coordination) or a broadband increase in aperiodic offset (as a proxy of changes in baseline neuronal spiking), two distinct mechanisms that absolute power could not dissociate. These different approaches may explain some inconsistencies in the developmental literature when oscillatory frequencies are not determined per infant (e.g. reports of both decreasing (Whitehead et al., 2018) and increasing (Yrjölä et al., 2024) low delta band power from the late preterm period towards TEA in active sleep). Separating oscillatory and non-oscillatory activity could therefore allow a more specific characterization of developmental changes by clarifying the respective contribution of background activity and oscillations to the observed power changes throughout development.

A few recent studies using spectral parameterization have begun to examine developmental changes in the provided spectral metrics. Aperiodic offset shows an increase from 2 months to 1 year of age (Wilkinson et al., 2024) and similarly from 6 to 16 months (Rico-Picó et al., 2023). Aperiodic exponent decreases from childhood into adolescence/adulthood (Favaro et al., 2023; Hill et al., 2022), but findings in infancy are scarce and indicate a less clear picture, as noted in the systematic review of Stanyard et al., (2024). Within infancy, the exponent is shown to increase between 2 and 8 months with slower changes thereafter (Wilkinson et al., 2024). A study even reported a decrease between 6 and 16 months of age (Rico-Picó et al., 2023), when broad frequency ranges were analyzed (but see Schaworonkow et al., 2021 for narrow-band characterization). Periodic activity also undergoes marked age-related changes, indicating the gradual emergence and differential trajectories for distinct oscillations along development (Li, et al., 2025.; Wilkinson et al., 2024). Among these, theta activity emerges particularly early in brain development, and periodic theta power mostly increases from 2 months of age towards the end of the first postnatal year, before subsequently declining (Rico-Picó et al., 2023; Wilkinson et al., 2024). These findings in early development have also underlined notable regional differences particularly along the posterior-anterior axis, with posterior regions showing larger offset values in the first year as well as lower theta power after 9-12 months of age compared to anterior regions (Rico-Picó et al., 2023; Wilkinson et al., 2024).

Only a few studies have quantified the early developmental trajectory of parameterized spectral metrics in the neonatal period. Existing evidence indicates an increase in aperiodic exponent between 35-45 post-conception weeks (Chini et al., 2022; Luotonen et al., 2025), with single time-point assessments suggesting that aperiodic measures may serve as early markers of risk for neurodevelopmental atypicalities such as autism spectrum disorders or attention deficit and hyperactivity disorder (Karalunas et al., 2022; Shuffrey et al., 2022). However, to our knowledge, no work to date has tracked the early longitudinal developmental evolution of these measures in preterm infants, in the early post-term period, and in comparison with full-term neonates. This gap applies to both aperiodic and periodic metrics, and addressing it would refine our understanding of traditional power spectral measures through a better delineation of background neural activity and oscillations such as theta. Characterizing the evolution of aperiodic and periodic brain activity during the post-term period might further highlight some clinically-relevant variability across infants, which needs to be investigated.

Moreover, the potential relationships between EEG aperiodic/periodic features and the brain anatomical characteristics and maturation have never been explored in the early developmental period. Multi-modal magnetic resonance imaging (MRI) can provide complementary insights into neonatal and infant brain structure (Dubois et al., 2021) and it is widely used to assess brain lesions, abnormalities and/or dysmaturation processes in preterm infants. In particular, the microstructure of brain tissues can be assessed with diffusion MRI, providing markers that are mostly sensitive to the cell and membrane density, the dendritic arborization in the gray matter, and to the fiber organization and myelination in the white matter (Ouyang et al., 2019). Exploring whether aperiodic features and brain oscillations are differently sensitive to these microstructural properties during early development might help understand their ontogeny and their role in the setting up of cerebral networks.

In this study, we aimed to investigate the early maturation of the brain activity in infants born very premature compared to full-terms, using high-density EEG recordings at TEA and 2 months later, and a spectral parameterization approach to dissociate between aperiodic and periodic activities. We focused on brain activity during active/REM sleep because it is the most prevalent sleep state in neonates (Grigg-Damberger, 2016); it plays a crucial role in supporting the development of functional networks (Del Rio-Bermudez & Blumberg, 2018), and it allows measuring robust oscillations such as theta, a rhythm sensitive to early vulnerability associated with prematurity (Tokariev et al., 2019). In particular, we aimed to investigate how prematurity impacts early brain functioning by: (i) examining how the spectral profiles evolve from TEA to 2 months post term, (ii) comparing periodic and aperiodic profiles between preterm and full-term infants, and (iii) exploring how clinical risk factors and the microstructural maturation of brain tissues account for the inter-individual variability within the preterm group at TEA. More specifically, we hypothesized that developmental changes between TEA and 2 months would be reflected in an increase in aperiodic offset, given the substantial increase in absolute power characterizing this developmental period, and in an increase in theta activity power based on prior developmental observations. Besides, we hypothesized that preterm infants may show altered aperiodic and periodic features compared to full-term infants. Finally, we anticipated that the variability observed across preterms at TEA might be partly explained by differences in clinical characteristics and in the brain microstructural maturation.

## 2. Methods

### 2.1 Subjects and EEG recordings

This study included preterm infants who were admitted to Robert-Debré Children’s Hospital (Paris, France) in the Neonatal Intensive Care Unit (NICU) closely after birth and who benefitted from clinical MRI at TEA, as well as full-term neonates without any neurodevelopmental risk factors (Adibpour et al., 2025). Infants were scheduled for high-density EEG recordings at two longitudinal timepoints: first, at TEA for preterms or close to birth for full-terms, and secondly two months later (corresponding to 2mCA for preterms). The study protocol was approved by the ethics committee (Comité de Protection des Personnes, CPP Ile de France 3), and parents were informed about the procedure and gave written informed consent.

#### 2.1.1 Preterm group

From an initial group of 43 infants born before 32 weeks GA, without moderate-to-severe brain anomaly confirmed on MRI at TEA with Kidokoro scoring (Elbaz et al., 2026), this study considered 41 infants (mean and standard deviation in GA at birth: 26.9 ± 1.7 weeks, range: [24.1;30.8] weeks; 22 males, 19 females) who had adequate EEG recordings in active/REM sleep at least at one age (see criteria below). Data were obtained for 40 infants for the first assessment at TEA (postmenstrual age (PMA): 40.9 ± 0.8 [38.4;42.1] weeks; postnatal age: 13.9 ± 2.0 [10.0;17.6] weeks; 22 males, 18 females), and for 16 infants for the second EEG assessment at 2mCA (PMA: 50.2 ± 0.5 [49.1;51.0] weeks; postnatal age: 23.2 ± 1.7 [19.2;25.8] weeks; 9 males, 7 females). The remaining 27 infants were not included in the analysis due to attrition (n=13), participation in an intervention program between TEA and 2mCA (n=8), or insufficient duration of high-quality recording in active/REM sleep (n=6). In total, 15 preterm infants contributed with longitudinal EEG data at both TEA and 2mCA.

For preterm infants with EEG data at TEA, we considered some clinical characteristics that are well established risk factors for adverse outcomes in preterm populations (Pierrat et al., 2021; Sacchi et al., 2020; Vliegenthart et al., 2019), and that we have shown to influence brain measures derived from EEG (Adibpour et al., 2025) and MRI (Elbaz et al., 2026) in preterm infants. Specifically, we considered male sex, GA at birth, being small for GA at birth (defined as birth weight < 10th percentile for GA), and invasive mechanical ventilation lasting more than 1 day. Consistent with our previous works (Adibpour et al., 2025; Elbaz et al., 2026), we categorized GA at birth into three groups: Group I (GI: 24 weeks ≤ GA ≤ 26 weeks), Group II (GII: 26 weeks < GA ≤ 28 weeks), and Group III (GIII: 28 weeks < GA ≤ 32 weeks). Table 1 summarizes clinical and demographic characteristics of preterm infants with EEG data at TEA, as well as infants with longitudinal data at TEA and 2mCA. Clinical characteristics of infants with and without longitudinal data were comparable (see Sup. Table 1).

**Table 1.** Demographic and clinical characteristics of the preterm cohort at TEA (N = 40) and longitudinal subset (n = 15)

| Parameter | Group I<br>(n = 13) | Group II<br>(n = 17) | Group III<br>(n = 10) |
| --- | --- | --- | --- |
| Sex Male/Female, n(%) | 6/7 (46.2/53.8%) | 11/6 (64.7/35.3%) | 5/5 (50/50 %) |
| Small weight for GA at birth, n(%) | 2 (15.4%) | 1 (5.9%) | 4 (40.0%) |
| Invasive ventilation >1 day, n(%) | 7 (53.8%) | 4 (23.5%) | 5 (50.0%) |
| Longitudinal subset | n = 4 | n = 8 | n = 3 |
| Sex Male/Female, n(%) | 1/3 (25.0/75.0%) | 6/2 (75.0/25.0%) | 1/2 (33.3/66.7%) |
| Small weight for GA at birth, n(%) | 1 (25.0%) | 0 (0.0%) | 1 (33.3%) |
| Invasive ventilation >1 day, n(%) | 4 (100.0%) | 1 (12.5%) | 1 (33.3%) |

Abbreviations: GA, gestational age; TEA, term-equivalent age
Groups defined by GA at birth: Group I, $24+0 \leq GA \leq 26+0$ weeks; Group II, $26+1 \leq GA \leq 28+0$ weeks; Group III, $28+1 \leq GA \leq 32+0$ weeks. The longitudinal subset comprises infants with EEG recordings at both time points. Percentages are computed within each group relative to that group's own n.

#### 2.1.2 Full-term group

From an initial group of 16 neonates born at term and tested with EEG, this study considered 13 neonates (GA at birth: 40.1 ± 1 [37.6;41.6] weeks; 7 males, 6 females), with adequate EEG recording in active/REM sleep at least at one age. Data were obtained for 11 infants at the first assessment, a few days after birth so equivalent to TEA (PMA: 41.8 ± 1 [39.7;43.3] weeks; postnatal age: 1.8 ± 0.7 [0.6;2.9] weeks; 6 males, 5 females), and for 10 infants 2 months later (PMA: 50.3 ± 0.9 [49.3;51.7] weeks; postnatal age: 10.3 ± 1.6 [8.4;13.1] weeks; 5 males, 5 females). Among these, 8 infants had longitudinal data.

#### 2.1.3 EEG Recordings

Brain activity was recorded with a 128-channel high-density EEG system (HydroCel Geodesic Sensor Net; Magstim EGI, USA), with net size adapted to the infant-head circumference. Signal was digitalized at a sampling rate of 1000 Hz. Recordings were obtained while infants were naturally sleeping in a cot or occasionally in parents’ arms when this facilitated calming the baby. A synchronized camera recorded infant behavior to inform subsequent sleep-stage classification.

### 2.2. EEG Processing

#### 2.2.1 Sleep Scoring

Infant vigilance states were classified through a two-step process that is detailed in Adibpour et al. (2025). Initially, during the recording, the experimenter (P.A. or A.P.) marked the infant’s vigilance state based on behavior (eye opening and body movements). These annotations were then refined offline by a neonatal neurophysiology expert (H.N). This study focused on active sleep and its presumed developmental equivalent at 2mCA, REM sleep which is characterized by low-amplitude irregular and mixed frequency EEG activity predominantly within the theta and delta ranges (*AASM Sleep Scoring and Event Manual*, 2023.; Grigg-Damberger, 2016).

#### 2.2.2 Preprocessing

Preprocessing was performed in EEGLAB (Delorme & Makeig, 2004), as detailed in Adibpour et al., (2025). In brief, data were initially filtered between (0.2-45 Hz), down-sampled to 250 Hz and preprocessed using the Automated Pipeline for Infants Continuous EEG (APICE) (Fló et al., 2022). Artifact detection and correction involved the identification and interpolation of persistently noisy channels, transient artifacts, as well as removal of motion-related contaminations. Following automated preprocessing, all recordings underwent visual inspection to manually flag residual artifacted periods missed by the pipeline. Finally, data were re-referenced to the common average reference.

Following sleep staging and EEG preprocessing, artifact-free segments were included. Only infants with at least 1 minute of sufficient quality data were then included in the study.

#### 2.2.3 Computation of EEG Power Spectrum

The power spectral density (PSD) was computed using Welch’s method implemented in MATLAB, using the EEGLAB (Delorme & Makeig, 2004) and FieldTrip (Oostenveld et al., 2011) toolboxes. For each channel, the power spectrum was estimated using a window length of 8 seconds with 50% overlap. Analyses focused on the [0.5-30] Hz frequency range, which is relevant for neonatal EEG and consistent with previous spectral analyses in newborns (Fransson et al., 2013; Karalunas et al., 2022; O’Toole & Boylan, 2019). We used frequency bins of 0.125 Hz width. The absolute power values were converted to an equivalence of decibel scale by taking the base-10 logarithm of the power. The PSD was averaged across all EEG channels for each infant, to obtain a single whole-scalp power spectrum per subject.

### 2.3 Analyses of aperiodic and periodic components

#### 2.3.1 Parameterization of the power spectrum into Aperiodic and Periodic Components

Power spectral densities were decomposed into their aperiodic (1/f) and periodic (oscillatory) components using the FOOOF (Fitting Oscillations & One Over F) algorithm implemented in Python (Donoghue, Haller, et al., 2020), and ran from MATLAB via the py.fooof interface (Figure 1). This algorithm models the log-transformed power spectrum *P* at each frequency *f*, as the sum of an aperiodic background component and a set of Gaussian-distributed periodic peaks superimposed on that background. The model is described as:

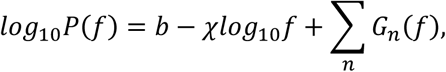

where *P*(*f*) is the absolute power at frequency *f*, *b* is the offset of the aperiodic component, corresponding to the intercept of the log transformed power spectrum, and *χ* is the exponent of aperiodic component, reflecting the slope of spectral decay (Figure 1), with larger exponent values indicating a steeper drop in power from low to high frequencies. The term *G_n_*(*f*) represents the Gaussian functions modeling the periodic components and is described as:

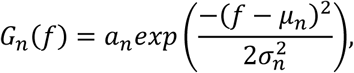

where *a_n_* is the peak amplitude, *μ_n_* is the center frequency, and *σ_n_* is the standard deviation, corresponding to the bandwidth of the Gaussian (Donoghue, Haller, et al., 2020).

**Figure 1.**
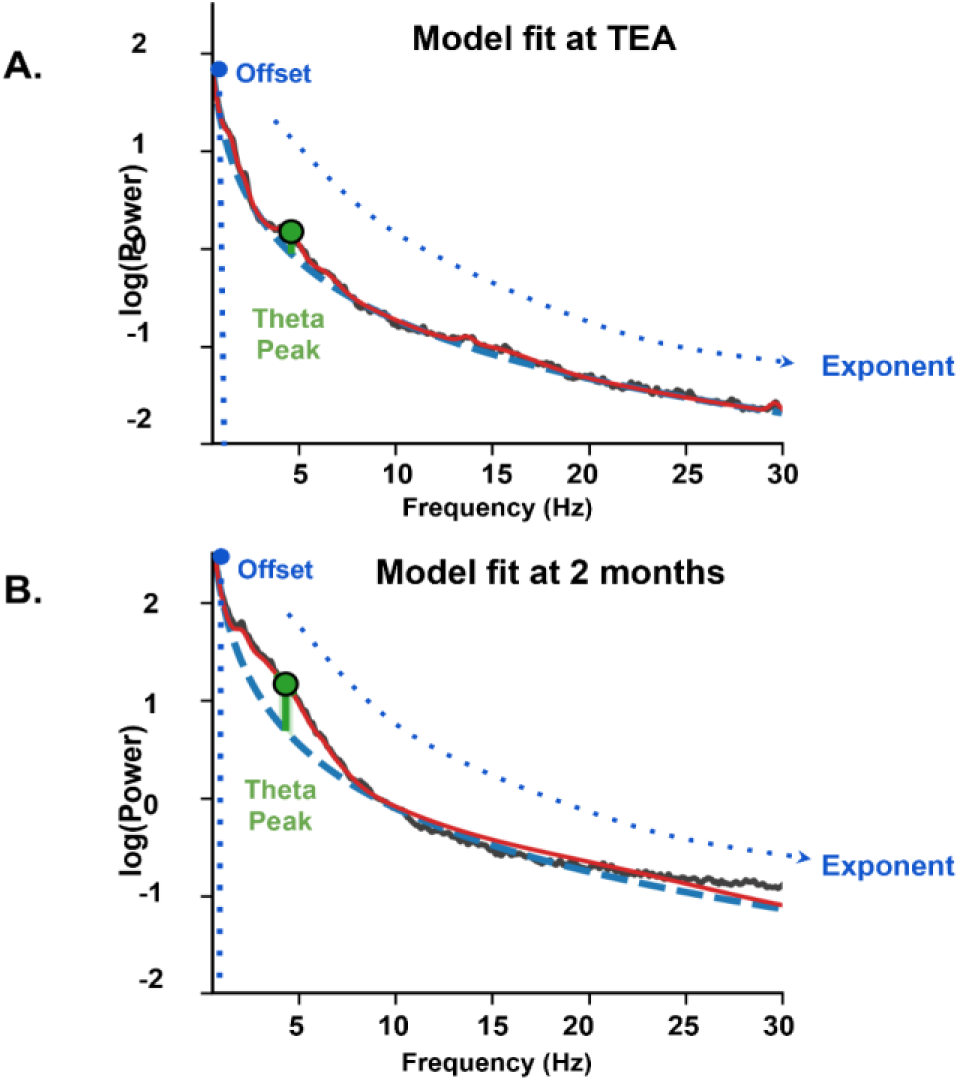
Illustration of aperiodic and periodic components. Two examples of the individual FOOOF model fit (red line) of the absolute power (black line) from (A) a preterm infant at TEA, and (B) a full-term infant at 2 months, respectively. Power spectrum is decomposed into aperiodic (dashed blue line) and periodic components, indicating the presence of a clear peak (solid green circle) in the theta range (4-7 Hz) above the aperiodic fit. Dotted blue lines represent aperiodic exponent and offset.

#### 2.3.2 Settings and extraction of metrics of interest

Spectral parameterization was first applied over the 0.5-30 Hz frequency range (Model #1). The aperiodic component was fitted and subtracted using the “fixed mode”, as infant spectra typically lack the clear low-frequency inflection point (“knee”) observed in adults (Cellier et al., 2021; Donoghue, Haller, et al., 2020; Wilkinson et al., 2024). To estimate the periodic components, a maximum of 7 peaks with width limits set to 0.5-18 Hz were considered to ensure better fitting performance for infant recordings, consistent with a prior developmental EEG study (Wilkinson et al., 2024). For each infant, metrics of offset and exponent were extracted for the aperiodic component in the fitted 0.5-30 Hz model (see goodness of fit in Sup. Table 2). For the periodic component, theta power was extracted by focusing on the 4-7Hz range, a range where rhythmic oscillatory activity is observed at 0-2 months of age (Grigg-Damberger, 2016; Wallois et al., 2021), and particularly dominate during active/REM sleep in neonates and infants (GriggDamberger, 2016). The initial model fit (0.5-30 Hz) for the periodic components failed to reliably capture a distinct theta peak for some infants (see Sup. Table 3, 4), primarily due to increased spectral noise at higher frequencies, which can obscure estimations of periodic components, an issue previously noted in infant EEG studies (Schaworonkow & Voytek, 2021). For the estimation of the periodic activity parameters, the spectrum was therefore parameterized over a narrower frequency range of 0.5-10 Hz (Model #2), while retaining all other model settings. This narrower range improved the precision of theta peak detection by limiting the influence of high-frequency noise (Schaworonkow & Voytek, 2021) and preserved data from infants whose theta peak were not reliably captured under Model #1, thereby maximizing the usable sample size. Within the theta range, we extracted the log-transformed power of the most prominent identified peak as the measure of periodic power.

**Table 2.**
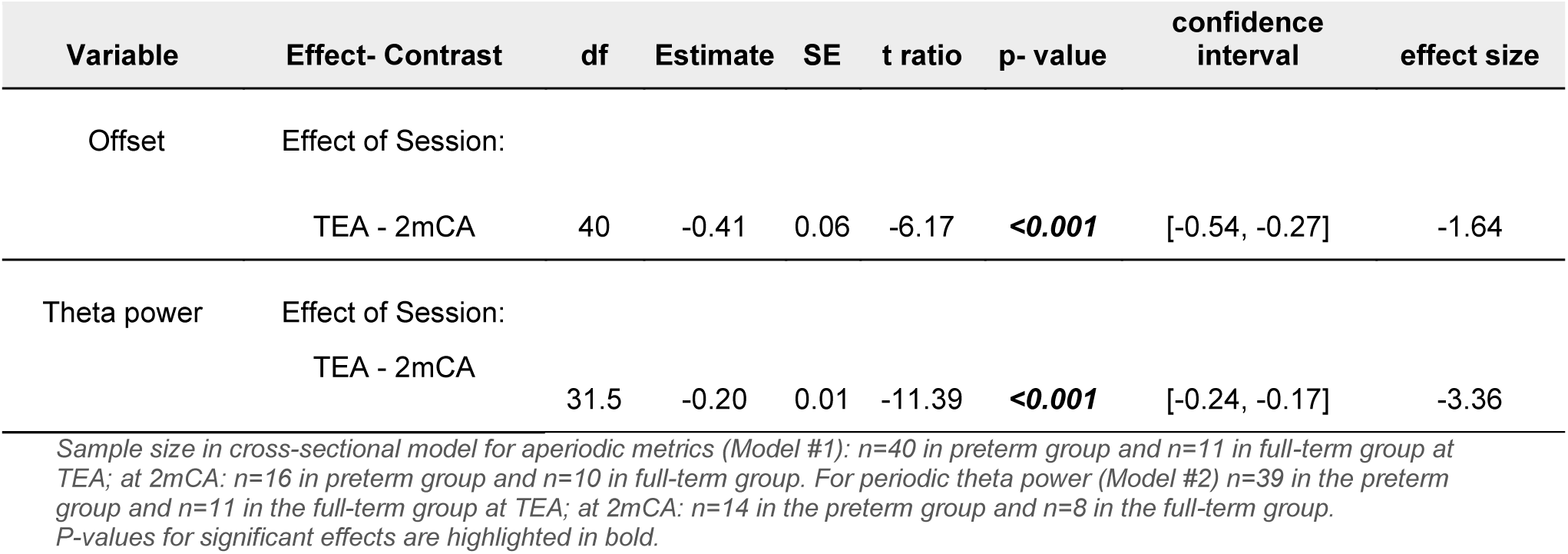
Summary of post-hoc tests for the significant effects of models on global measures (cross-sectional cohort)

| Variable | Effect- Contrast | df | Estimate | SE | t ratio | p- value | confidence interval | effect size |
| --- | --- | --- | --- | --- | --- | --- | --- | --- |
| Offset | Effect of Session: |  |  |  |  |  |  |  |
|  | TEA - 2mCA | 40 | -0.41 | 0.06 | -6.17 | **<0.001** | [-0.54, -0.27] | -1.64 |
| Theta power | Effect of Session: |  |  |  |  |  |  |  |
|  | TEA - 2mCA | 31.5 | -0.20 | 0.01 | -11.39 | **<0.001** | [-0.24, -0.17] | -3.36 |

Sample size in cross-sectional model for aperiodic metrics (Model #1): $n=40$ in preterm group and $n=11$ in full-term group at TEA; at 2mCA: $n=16$ in preterm group and $n=10$ in full-term group. For periodic theta power (Model #2) $n=39$ in the preterm group and $n=11$ in the full-term group at TEA; at 2mCA: $n=14$ in the preterm group and $n=8$ in the full-term group. P-values for significant effects are highlighted in bold.

**Table 3.**
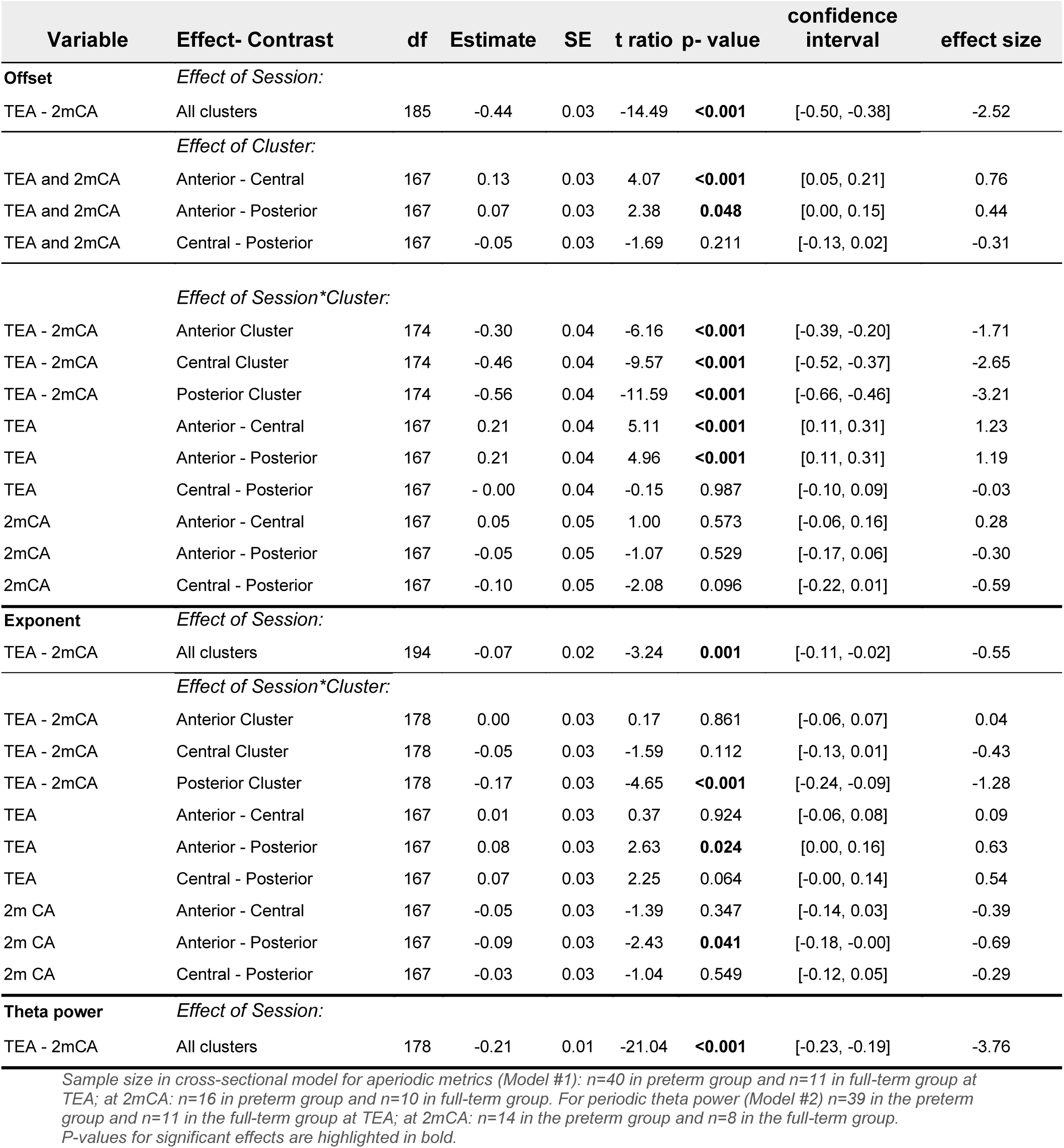
Summary of post-hoc tests for the significant effects of models on cluster measures (cross-sectional cohort)

Note that Sup. Tables 2-4 present validation analyses on model fit and periodic theta estimations extracted with Model #1 and Model #2, for the subgroup of infants in whom theta peak was reliably captured in Model #1.

**Table 4.** Summary of post-hoc tests for the significant effects of clinical risk factors on TEA measures.

| Variable | Effect- Contrast | df | Estimate | SE | t ratio | p- value | confidence interval | effect size |
| --- | --- | --- | --- | --- | --- | --- | --- | --- |
| Offset | **Effect of invasive ventilation** |  |  |  |  |  |  |  |
|  | yes - no | 34 | -0.25 | 0.08 | -2.83 | **0.007** | [-0.43, -0.07] | -0.97 |
| Exponent | **Effect of GA at birth** |  |  |  |  |  |  |  |
|  | Group II - Group III | 34 | -0.18 | 0.06 | -2.86 | **0.013** | [-0.33, -0.03] | -0.98 |
|  | Group I - Group III | 34 | -0.18 | 0.06 | -2.89 | **0.012** | [-0.33, -0.03] | -0.99 |
|  | **Effect of invasive ventilation** |  |  |  |  |  |  |  |
|  | yes - no | 34 | -0.13 | 0.05 | -2.52 | **0.016** | [-0.23, -0.02] | -0.86 |
| Theta power | **Effect of sex** |  |  |  |  |  |  |  |
|  | male - female | 33 | -0.05 | 0.01 | -2.68 | **0.01** | [-0.08, -0.01] | -0.93 |
|  | **Effect of small weight for GA** |  |  |  |  |  |  |  |
|  | yes - no | 33 | 0.05 | 0.02 | 2.31 | **0.02** | [0.00, 0.10] | 0.80 |

Sample size of preterm group at TEA $n = 40$ . P-values for significant effects are highlighted in bold.

#### 2.3.3 Computation over spatial clusters

Because maturation proceeds differently across brain regions during early development (Dubois et al., 2014), averaging EEG metrics over all electrodes and thus over the whole brain might obscure developmental effects. Thus, we examined whether extracting metrics over more localized electrode clusters would reveal distinct maturational profiles across brain regions, as suggested by previous developmental studies showing spatial differences in absolute power and in aperiodic and periodic components particularly along the posterior-anterior axis (Novelli et al., 2016; Rico-Picó et al., 2023; Schaworonkow & Voytek, 2021; Stanyard et al., 2024). To account for this spatial heterogeneity, we computed EEG metrics separately for three bilateral clusters: frontal (36 electrodes), central (36 electrodes), and posterior (37 electrodes), following the parcellation scheme used by (Calbi et al., 2019) for the 128-channels Hydrocel Geodesic Sensor Net.

As for the global set of electrodes described in 2.3.2, spectral parameterization was performed for each cluster of electrodes independently. Model #1 was used to extract the aperiodic metrics (offset and exponent) and Model #2 to extract the periodic theta metrics.

### 2.4 Collection and analyses of MRI data in preterm infants at TEA

In addition to EEG assessments, preterm infants had a clinical MRI at TEA either because they were born extremely premature (before 28 weeks GA), or because of suspicion of cerebral anomalies. TEA-MRI was performed on a 3T scanner on the same day as EEG for most infants (except for 3, for whom the two exams were separated by no more than 6 days). A standardized protocol was used (see details in Elbaz et al, 2026, Devisscher et al, 2026), including T2-weighted sequences in axial, coronal and sagittal planes (echo time [TE] = 150 ms, repetition time [TR] = 5500 ms), 3D-T1-weighted and susceptibility-weighted sequences. A diffusion-weighted sequence with 42 directions at b=1000s/mm^2^ (TE = 75 ms, TR = 5000 ms, 2mm-isotropic resolution) was also acquired. First, radiological examinations with scoring adapted to the preterm population (Kidokoro et al., 2013; Elbaz et al., 2026) indicated that all infants included in this EEG study had normal MRI or mild abnormalities (whole-brain Kidokoro scores ≤ 7). Second, pre- and post-processing of images were performed. T2-weighted sequences of different planes were combined with a super-resolution algorithm to obtain a 0.8 mm-isotropic spatial resolution (Ebner et al., 2020). Brain tissues were segmented to provide masks of the cortex and white matter (Elbaz et al., 2026). Diffusion-weighted images were post-processed to correct for motion artefacts and susceptibility-induced distortions (Devisscher et al, 2026). The model of diffusion tensor imaging (DTI) was estimated, and quantitative maps of fractional anisotropy (FA) and mean diffusivity (MD) were generated. The T2-weighted and b=0 images were non-linearly registered (Devisscher et al, 2026), enabling the projection of brain tissue masks on DTI maps (Figure 2). Averaged FA and MD metrics were then computed over the masks of cortex and white matter.

**Figure 2.**
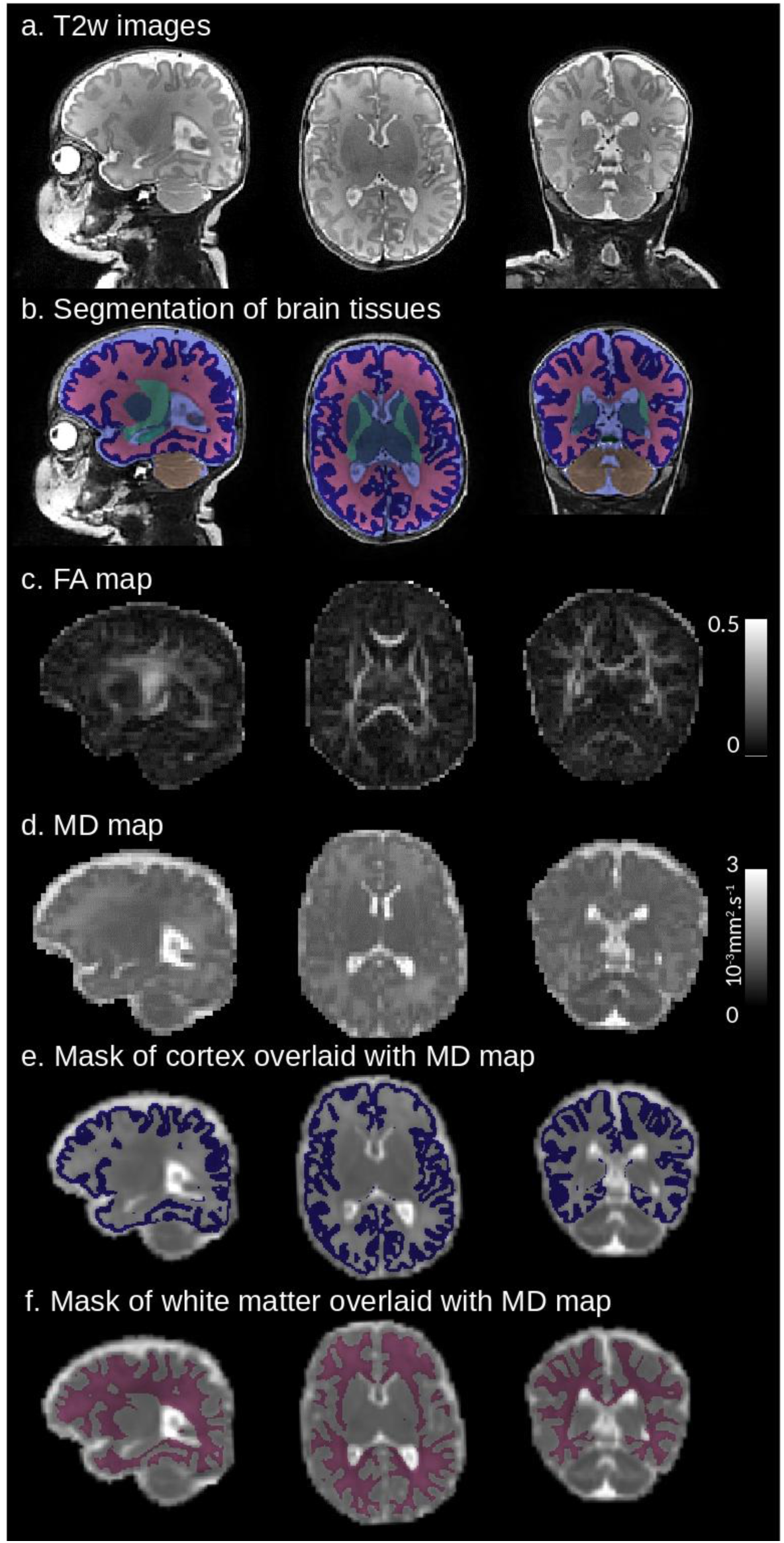
Multimodal MRI images of a preterm infant at term-equivalent age. T2-weighted (T2w) images after super-resolution reconstruction are shown on sagittal, axial and coronal slices (a). These enabled the segmentation of brain tissues (b) in particular the cortex (dark blue) and the white matter (pink). Maps derived from diffusion tensor imaging (DTI) are shown at similar levels of the brain: fractional anisotropy (FA) (c) and mean diffusivity (MD) (d), onto which masks of the cortex (e) and white matter (f) have been superimposed following non-linear registration between T2w images and b=0 images.

### 2.5 Statistical analyses

#### 2.5.1 Analysis of maturation and prematurity effects

To investigate whether EEG-derived metrics evolve between TEA and 2mCA, and differ between the preterm and full-term groups, we employed linear mixed-effects models (LMM) on the entire dataset (both cross-sectional and longitudinal data of the two groups, n=54; 41 preterms + 13 full-terms). The dependent variable was either aperiodic offset, aperiodic exponent, or theta power. Group (preterm vs. full-term), Session (TEA vs. 2mCA) and their interaction were considered as fixed-effect predictors, and subjects as random-effect predictors. When a significant main effect or interaction was observed, post-hoc tests were conducted using pairwise comparisons of estimated marginal means. All models were fitted in R using the lme4 package (Bates et al., 2015), with p-values for fixed effects obtained with Satterthwaite’s approximation of degrees of freedom (lmerTest).

To isolate genuine developmental change, we further estimated an LMM model only on infants with longitudinal data from both sessions (n = 23; 15 preterms + 8 full-terms).

#### 2.5.2 Analysis of spatial-heterogeneity effects

In line with prior analyses, LMM models were fitted for metrics estimated over each spatial cluster, in the entire dataset (n=54). In addition to the effects of Group (preterm vs. full-term) and Session (TEA vs. 2mCA), the effect of Cluster (Anterior, Central, Posterior) and their interactions were included as fixed-effect predictors and subjects as random-effect predictors. P-values for fixed-effects were extracted using Type III analysis of variance (ANOVA) F-tests with Satterthwaite’s approximation of degrees of freedom. This approach allowed appropriate estimation of predictors with more than two levels (i.e. three-level Cluster variable) and their interactions with other predictors, after accounting for all other terms in the model. Post-hoc comparisons of estimated marginal means were conducted for significant effects.

#### 2.5.3 Analysis of the effects of clinical factors and brain microstructural maturation in preterms at TEA

We further aimed to investigate the factors that might explain the inter-individual variability of EEG metrics in preterm infants at TEA (n=40). For this, we focused on whole brain metrics, and used ANOVA with aperiodic offset, exponent, and theta-band power included as dependent variables. As independent variables, we considered the infant Sex (male/female), the group of GA at birth (GI, GII, GIII), small weight for GA at birth (yes/no), and history of invasive mechanical ventilation (yes/no) (Adibpour et al., 2025; Elbaz et al., 2026). Significant main effects were followed up with post-hoc analyses using estimated marginal means.

We also evaluated whether the whole brain EEG metrics were dependent on the microstructural maturation of brain tissues at TEA. For each EEG metric (aperiodic offset, aperiodic exponent, and theta power), we fitted linear regression models including a DTI metric (either FA or MD) averaged either over the white matter or the cortex as covariate, while adjusting for age at EEG/MRI acquisition.

Prior to statistical inference, we verified the standard assumptions of linear modeling through visual inspection of residual plots. These included checks for linearity, homoscedasticity, and normality of residuals. To assess multicollinearity among independent variables, we used the Variance Inflation Factor (VIF), with all VIF values < 3, indicating acceptable levels of collinearity. When multiple models were run, p-values were corrected for multiple comparisons with false discovery rate approach (FDR) and statistical significance was set at p < 0.05 after correction.

In a set of validation analyses, we also re-estimated all statistical models for periodic theta power in the subgroup of infants in whom theta peak was reliably captured in Model #1 (see Sup. Table 5).

## 3. Results

On average, the mean usable duration of active sleep at TEA was 6.14 ± 2.52 minutes (1.04-12.32 minutes) for preterm infants (n = 40), and 5.94 ± 2.26 minutes (2.52-9.61 minutes) for full-term neonates at birth (n = 11). At 2mCA, duration of usable REM data was 5. 65± 2.49 minutes (1.30-10.43 minutes) for preterm infants (n = 16), and 6.74 ± 4.47 minutes (1.24-14.39 minutes) for full-term infants (n = 10). Because recording duration varied across infants and sessions, we verified that it did not influence our spectral estimates. Bivariate correlations between duration and each FOOOF-derived global metric were non-significant at both timepoints (all |r| ≤ 0.17, all p ≥ 0.43; FDR-adjusted p ≥ 0.93). Recording duration was also included as a covariate in all linear mixed-effects models reported below for global metrics, and was a non-significant predictor in every case (all p ≥ 0.56).

### 3.1 Qualitative description of power spectrums and model fitting

The PSD estimates for preterm and full-term infants at TEA and 2mCA are presented in Figure 3. Power spectra showed the typical 1/f distribution (Figure 3A). Inspection of average PSD at the global level revealed strong differences between TEA and 2mCA in both groups, with higher power across all frequencies in infants at 2mCA than at TEA, particularly within the 2- 7 Hz range. These differences in spectral profiles between TEA and 2mCA were also apparent in each cluster but appeared less pronounced in the anterior cluster than in the central and posterior ones (Figure 3A). Inspection of topographical distribution of PSD within theta band (4-7 Hz), indicated qualitative differences between the preterm and full-term groups in particular for the contrast between 2mCA/2m and TEA/0m (Figure 3B-C).

**Figure 3.**
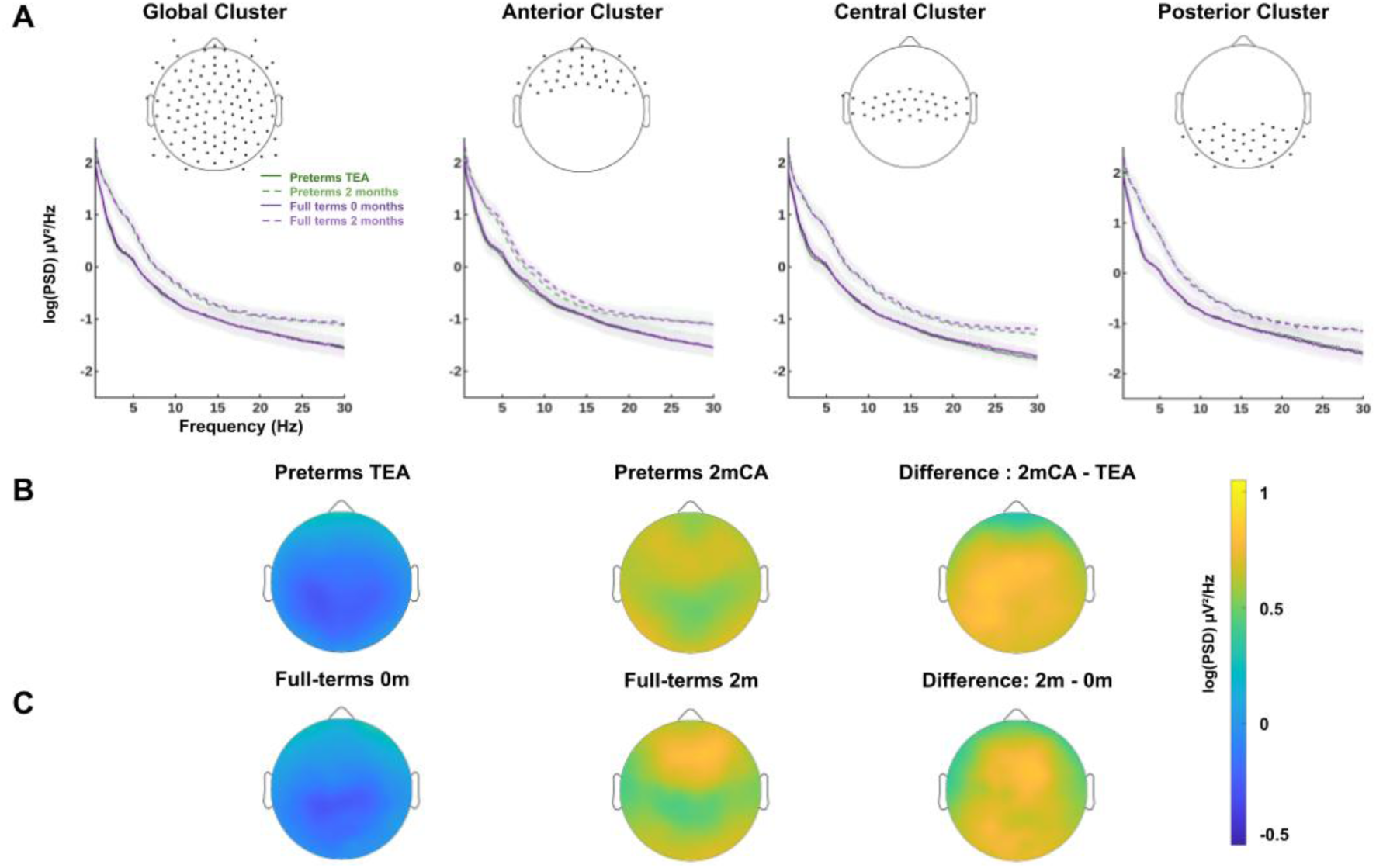
Group-averaged power spectral density (PSD) and topographies across preterm and full-term infants. (A) Group-averaged log-transformed PSD (µV²/Hz) computed across the 0.5-30 Hz range for preterms at TEA (solid green), preterms at 2mCA (dashed green), full-terms at 0 months (solid purple), and full-terms at 2 months (dashed purple), shown for the global cluster and for the anterior, central, and posterior regional clusters. Shaded areas represent the standard error of the mean. Electrodes in each cluster are represented on top. (B) Topographical distribution of log(PSD) averaged across the 4-7 Hz range (theta activity) in preterm infants at TEA (left), at 2mCA (middle), and the difference map (2mCA-TEA, right). (C) Same topographical representation for full-term infants at 0 months (left), 2 months (middle), and the difference map (2m-0m, right).

For all infants, aperiodic metrics (offset and exponent) were extracted exclusively from Model #1, and theta power from Model #2 (see goodness of fit from both models in Sup. Table 2). Yet, with this latter model, no theta peak was detected for a few infants: at the global level (1 preterm at TEA, 2 preterms at 2mCA, 2 full-terms at 2m) and at the cluster level (anterior: 1 preterm at TEA, 1 preterm at 2mCA; central: 1 preterm at TEA, 2 preterms at 2mCA, posterior: 1 preterm at TEA, 2 preterms at 2mCA, 2 full-terms at 2m). Accordingly, analyses of periodic theta power in preterm infants at TEA (reported in sections 3.4 and 3.5) are based on n = 39, whereas analyses of aperiodic metrics use the full preterm sample (n = 40); analogous adjustments apply at 2mCA and for cluster-level analyses.

### 3.2 Effects of maturation and prematurity on aperiodic and periodic metrics

Aperiodic metrics and theta power averaged over all electrodes are presented in Figure 4 for preterm and full-term infants at TEA and 2mCA. Results of the LMM model performed on the global PSD estimations (Sup. Table 6) are summarized below, with corresponding post-hoc tests reported in Table 2. Analysis of aperiodic offset showed a significant main effect of Session (t(73) = 3.60, p < .001), with post-hoc tests indicating higher offset values at 2mCA compared to TEA. No significant effect was found for aperiodic exponent. Periodic theta power also showed a main effect of Session (t(34.07) = 6.91, p < .001), reflecting higher theta power at 2mCA compared to TEA (see Table 2 for post-hoc tests of these effects). Effects of Group or interactions were non-significant (all |t| < 1, p > .46).

**Figure 4.**
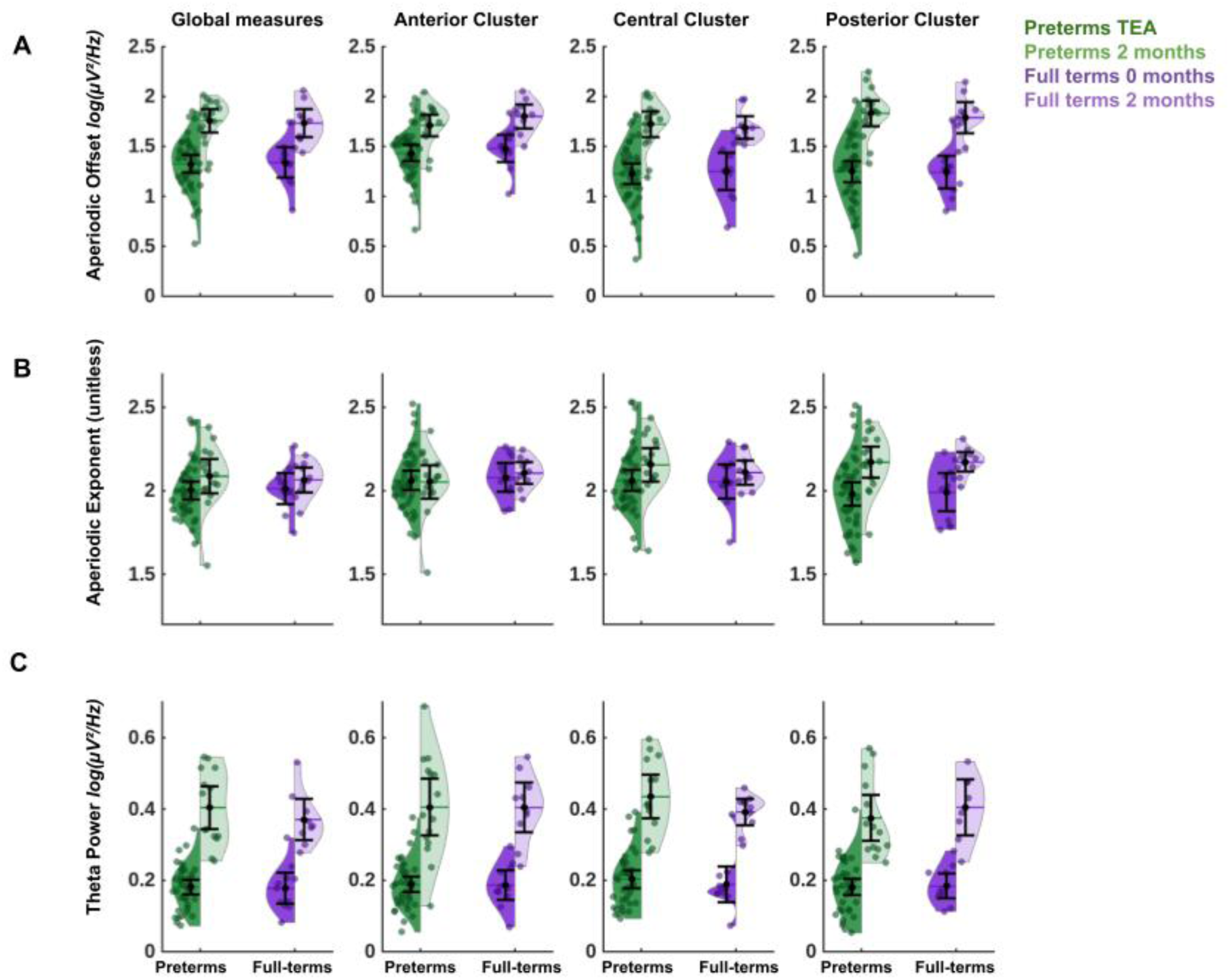
Group-averaged distribution of parameterized metrics across preterm and full-term infants. Violin plots show distributions of (A) aperiodic offset, (B) aperiodic exponent, and (C) theta power in preterms (green) and full-terms (purple) at TEA)/ 0 months and at 2mCA/ 2 months. Darker points represent individual infants. Offset and theta power show an effect of Session across all clusters, while exponent indicates a Session effect only in the posterior cluster, all highlighting an increase from TEA/0 months to 2months.

As a sanity check, we repeated these analyses in the subset of infants with longitudinal data (15 preterm and 8 full-term infants). The pattern of results remained consistent: i) aperiodic offset showed a significant main effect of Session (t(42) = 4.08, p < .001), with higher values at 2mCA compared to TEA, ii) no significant effect was found for aperiodic exponent, and iii) theta power showed a significant main effect of Session (t(18) = 6.49, p < .001), with higher power at 2mCA compared to TEA (see Sup. Table 7 for results of main model and Sup. Table 8 for post-hoc tests of these effects).

### 3.3 Effects of spatial localization on aperiodic and periodic metrics

For cluster-based PSD estimations, LMM results (Sup. Table 9) are summarized below, with follow-up post-hoc tests reported in Table 3. No effects of Group or Group × Session × Cluster interaction was observed for any metric (all F < 1.74, p > 0.18).

Analysis of aperiodic offset revealed a significant main effect of Session (F(1, 183.59) = 211.32, p < .0001), with post-hoc tests indicating higher offset values at 2mCA compared to TEA. It further showed a main effect of Cluster (F(2, 164.5) = 8.37, p < .001), with higher values in the anterior cluster compared to central and posterior clusters. A significant Session × Cluster interaction was also found (F(2, 164.5) = 8.29, p < .001), with post-hoc tests showing higher offset at 2mCA compared to TEA across all clusters, with the biggest effect size in the posterior cluster. This interaction also highlighted larger offset values over the anterior cluster compared to central and posterior cluster at TEA, but not at 2mCA.

Analysis of aperiodic exponent indicated a significant effect of Session (F(1, 192.3) = 10.64, p = 0.001), with post-hoc comparisons showing larger exponent values at 2mCA compared to TEA. A significant Session × Cluster interaction (F(2, 164.5) = 6.49 p < .001) further demonstrated higher exponent at 2mCA compared to TEA, in the posterior but not in the anterior or central clusters. This also highlighted larger exponent values at TEA over the anterior compared to the posterior cluster, whereas this pattern was reversed at 2mCA. Periodic theta power indicated a significant main effect of Session (F(1, 178.38) = 445.57, p < .001), reflecting higher power at 2mCA compared to TEA.

Analyses on infants with longitudinal data corroborated these observations (Sup. Tables 10, 11).

### 3.4 Effects of clinical factors on aperiodic metrics and theta power in preterms at TEA

We then focused on spectral metrics obtained in preterm infants at TEA. Results of the ANOVA models incorporating various risk factors (Sup. Table 12) are summarized below with post-hoc tests reported in Table 4.

Aperiodic offset revealed a significant effect of invasive ventilation (F(1, 34) = 6.95, p = .013), and post-hoc analyses indicated that infants who received ventilation exhibited lower offset values than those who had not been ventilated.

ANOVA for aperiodic exponent indicated a significant effect of GA at birth (F(2, 34) = 5.39, p = .009) with GIII group (i.e. higher GA at birth) presenting significantly higher exponent values compared to GII and GI. It also showed a significant effect of invasive ventilation (F(1, 34) = 6.38, p = .016), with post-hoc tests showing that infants who received ventilation had lower exponents than those who did not.

Periodic theta power further showed a significant effect of Sex (F(1, 33) = 4.68, p = 0.03) with lower theta power in males compared to females. It also showed a significant effect of being Small for GA at birth (F(1, 33) = 5.34, p = 0.02), surprisingly indicating that infants with small weight presented higher theta power compared to those without such risk factor.

### 3.5 Associations between brain microstructural maturation and EEG metrics at TEA

Finally, we examined whether spectral parameterized metrics in preterms at TEA were related to brain microstructure, indexed by FA or MD averaged over the cortex or the white matter. Linear models revealed no significant association for aperiodic metrics (offset, exponent: all |t| < 1.48, all p > 0.149). For theta power, no significant relationship was found with MD within cortical gray matter or white matter (all |t| < 0.63, all p > 0.53), nor with FA within white matter (t(34) = 0.58, p = 0.56). In contrast, theta power was significantly related to FA within cortical gray matter (t(34) = −3.11, p = 0.004), with higher theta power associated with lower cortical FA (Figure 5).

**Figure 5.**
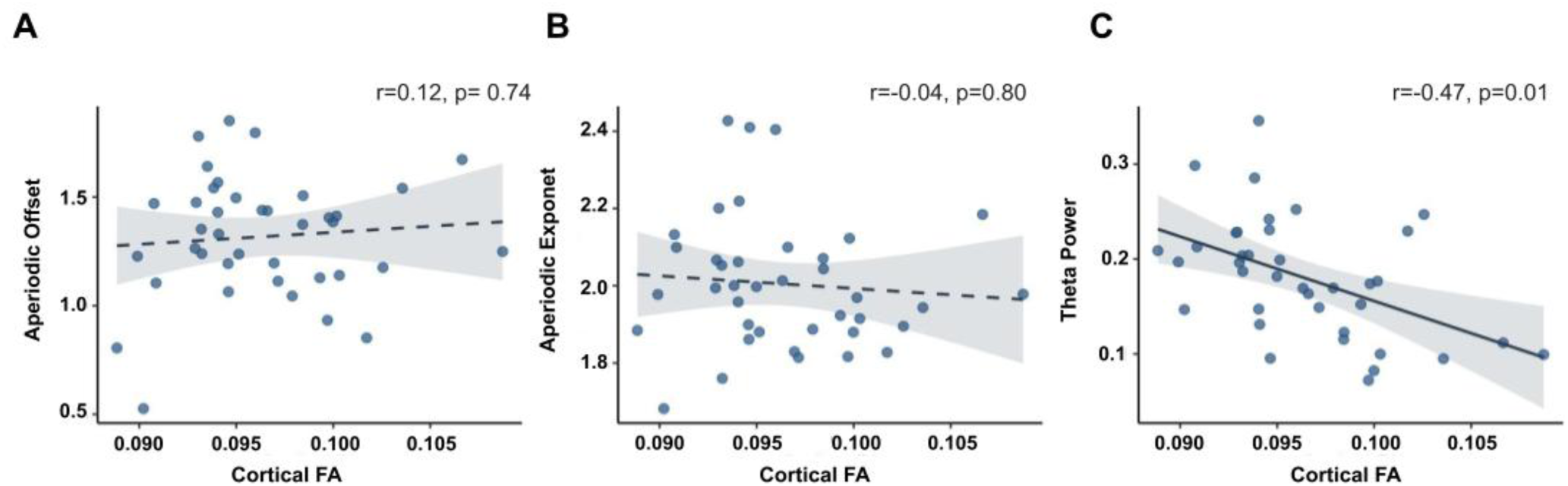
Associations between global EEG metrics and cortical fractional anisotropy (FA) at term-equivalent age in preterm infants. Scatter plots showing the relationship between cortical FA and (A) aperiodic offset, (B) aperiodic exponent, and (C) theta power, in preterm infants at TEA. Lines depict the predicted regression slope; shaded areas indicate 95% confidence intervals. Solid lines indicate statistically significant associations (p < 0.05), while dashed lines indicate non-significant ones. The partial correlation coefficients (r) and p-values displayed correspond to the effect of cortical FA in each model, with p-values FDR-corrected across the three outcomes.

## 4. Discussion

This study used high-density EEG recordings and spectral parameterization to characterize developmental changes in early brain activity from TEA to 2mCA in preterm and full-term infants. This approach allowed us to disentangle the evolution of the aperiodic background activity and periodic theta activity during active/REM sleep, shedding light on their respective contributions to the maturation of brain networks. Our analyses revealed robust age-related changes in aperiodic metrics and periodic theta power across this developmental window. Cluster-based analyses further uncovered region-specific developmental patterns that were undetectable by global measures, indicating that the maturation of brain activity progresses heterogeneously across different brain regions and networks. Although preterm and full-term infants exhibited broadly similar trajectories, they also showed substantial inter-individual variability. Within the preterm cohort, variability in spectral metrics at TEA was associated with distinct clinical risk factors and, to a lesser extent, with MRI-derived indices of cortical microstructural maturation.

### 4.1 Development of aperiodic and periodic activity during the early postnatal period

#### Age-related changes in aperiodic activity

Developmental trajectories of EEG aperiodic metrics have been documented in a few prior longitudinal investigations, targeting developmental windows after 1-2 months of age in typically developing infants (Rico-Picó et al., 2023; Schaworonkow & Voytek, 2021; Wilkinson et al., 2024) or the period between 35 and 46 weeks PMA (Chini et al., 2022). Our study extends this literature by characterizing the developmental period between TEA-2mCA, and by including insights from both preterm and full-term infants. Within this early window, we observed an increase in aperiodic offset, which appears to be an extension of previously reported age-related increases starting from 2 months towards one year of age (Rico-Picó et al., 2023; Wilkinson et al., 2024).

For aperiodic exponent, we also observed an age-related increase over posterior areas. This pattern aligns with previous reports of an increase between 35 and 46 weeks PMA (Chini et al., 2022), and between 2 and 8 months (Wilkinson et al., 2024), but contradicts a report of decrease from 1 to 7 months (Schaworonkow & Voytek, 2021). We suggest that this discrepancy likely stems from methodological differences and in particular the use of a restricted frequency range (1-10Hz) in the latter study, which may insufficiently represent the neural activity beyond the knee-point in the power spectra. This consideration applies specifically to the aperiodic exponent, where estimations of the slope of spectral decay benefits from a broader frequency range (Gerster et al., 2022). In the present study, all aperiodic estimates were derived from the broader 0.5-30 Hz fit (Model #1). The increase in aperiodic exponent is further supported by evidence from animal models within a broader fitting range (5-45 Hz), demonstrating a developmental increase across postnatal days 2-12 in mice medial prefrontal cortex (Chini et al., 2022).

The observation of age-related increases in aperiodic offset and exponent during infancy might appear counterintuitive given that these metrics show larger values in early infancy (Chini et al., 2022; Rico-Picó et al., 2023; Schaworonkow & Voytek, 2021) than in childhood and adolescence (up to 17 years old) (Favaro et al., 2023; Karalunas et al., 2022; Stanyard et al., 2024). But an integrated view rather suggests that the direction of changes in aperiodic metrics might reverse during development, with an inverted U-shaped trajectory. In particular, the exponent might decline after ∼6-8 months (Rico-Picó et al., 2023), while the offset might stabilize after ∼12 months (Rico-Picó et al., 2023; Wilkinson et al., 2024). Further longitudinal studies across the first postnatal year are needed to determine the precise timing of such inflection points.

The neurobiological mechanisms that drive the evolution of aperiodic activity in the early developmental period are not yet fully understood. However, the aperiodic offset, reflecting the overall EEG power, is thought to be a proxy for baseline neuronal spiking activity (Donoghue, Haller, et al., 2020; Manning et al., 2009). During the early postnatal period, dramatic increases in dendritic arborization and synaptogenesis occur (Huttenlocher & Dabholkar, 1997), which might cause such elevation in baseline neural activity. Reflecting the rate at which spectral power decays across frequencies, the aperiodic exponent has been linked to the balance between synaptic excitation (E) and inhibition (I) within cortical circuits (Donoghue, Haller, et al., 2020; Gao et al., 2017), with higher exponent associated with lower E/I ratio related to increased inhibitory activity (Gao et al., 2017). The observed increase in exponent from TEA to 2mCA therefore likely reflects ongoing maturation of excitatory and inhibitory networks. In particular, inhibitory interneurons continue migrating and integrating into cortical circuits, and GABAergic signaling transitions from excitatory to inhibitory (Kostović & Judaš, 2015; Peerboom & Wierenga, 2021; Xu et al., 2011). These processes enable the formation and strengthening of excitatory and inhibitory synapses which could change the E/I balance (Chini et al., 2022; Wilkinson et al., 2024), and contribute to the changes in aperiodic activity observed during early infancy.

#### Age-related changes in periodic activity

We further observed an increase in theta power in infants from TEA to 2mCA, which is consistent with previous studies showing an early modest increase from 2 months of age, followed by a decrease from the end of the first postnatal year (Rico-Picó et al., 2023; Wilkinson et al., 2024). Our findings highlighted that the increase in theta activity begins even earlier, at least from TEA.

Theta rhythm constitutes one of the earliest and most fundamental neural rhythms that is generated through interactions between glutamatergic and GABAergic neurons within basal forebrain nuclei and hippocampus (Buzsáki, 2002; Müller & Remy, 2018). Translational work has further demonstrated that theta activity mediates long-range connectivity, serving as a fundamental mechanism for integrating activity from distributed brain regions early on (Brockmann et al., 2011; Del Rio-Bermudez et al., 2017). In human development, theta is also suggested to play a crucial role in shaping large-scale networks, through a more precise synchronization between distant regions (Uhlhaas et al., 2010).

In keeping with this literature, the observed increase in theta power from TEA to 2mCA likely reflects the maturation of long-range connectivity, which sustain excitatory and inhibitory circuits and enable more precise rhythmic coordination across distributed cortical regions. This interpretation might be supported by our observation that theta power in preterm infants at TEA is partly related to the cortical maturation, as assessed with diffusion MRI. Cortical fractional anisotropy is known to decline during last gestational trimester in fetuses and preterm neonates, reflecting increasing microstructural complexity of the cortical plate as its initially radial organization, linked to apical dendrites and radial glia, is progressively reshaped by dendritic arborization and synaptogenesis (Ouyang et al., 2019). The inverse association we observed between theta power and cortical FA therefore suggests a parallel between increased oscillatory activity and progressive cortical maturation.

While this work focused on active/REM sleep and theta activity, future studies could perform a broader examination of oscillatory rhythms across sleep and wake states. In particular, delta activity during quiet/non-REM sleep may also serve as a sensitive marker of early maturation in preterm infants. However, accurate estimation of periodic delta activity remains challenging, as current spectral parameterization methods show limitations for model fitting at low frequencies, highlighting the need for models adapted to neonatal populations.

### 4.2 Spatial heterogeneity in early brain activity maturation

Early brain maturation is highly heterogeneous across regions during infancy, and our cluster level analyses were motivated by this in order to capture spatial differences in the development of early brain activity. We observed robust regional differences in aperiodic metrics at each age, as well as in the maturation rate of aperiodic metrics from TEA to 2 months, whereas periodic theta power was more spatially uniform. At TEA, anterior regions exhibited higher aperiodic offset and exponent values than posterior regions. By 2 months corrected age (2mCA), this spatial pattern had partially reversed: anterior regions no longer showed higher offset values, and the exponent was even greater in posterior than anterior areas. This redistribution coincided with regional differences in maturation rates. Specifically, increases in aperiodic offset were most pronounced in posterior areas relative to anterior and central areas, while the aperiodic exponent increased only in posterior areas, remaining stable elsewhere. In contrast, no significant regional differences were observed for periodic theta power, but further work considering more comprehensive regional analyses (i.e. beyond posterior-anterior directions) is needed to confirm a relative homogeneity of theta activity development across brain regions in this period.

The observation of higher aperiodic metrics in the anterior regions at TEA could appear misaligned to the classical posterior-to-anterior maturational gradient described during infancy. Yet, two considerations help reconcile our observations with this gradient. First, a maturational interpretation can only be drawn from the change between TEA and 2mCA. From this perspective, our findings align with the established earlier maturation of posterior areas, since the aperiodic exponent increased exclusively in posterior regions, and the offset likewise showed its largest increase in posterior areas. Second, the cross-sectional pattern at TEA is difficult to interpret, as this was the earliest measure in our cohort and the direction of preceding changes up to this time point was unknown. Higher offset and exponent values at TEA in anterior than posterior clusters may indicate a regional configuration shaped by several processes specific to this early postnatal window. Transient subplate neurons, for instance, are most numerous and persist longest in associative and prefrontal areas during this period (Kostović et al., 2015; Kostović & Judaš, 2010), potentially sustaining the rich background activity detected frontally at TEA. Another possibility is that immature GABAergic signaling before a shift from excitatory to inhibitory neurotransmission (Peerboom & Wierenga, 2021; Xu et al., 2011) may give rise to higher aperiodic activity due to the prevalence of excitatory and less mature inhibitory circuitries (Gao et al., 2017). These observations align with recent network modeling frameworks proposing that between 33 and 45 weeks of age, frontal regions lead the emergence of a synergistic scaffold in the neonatal brain, while sensory areas mature progressively from posterior to central regions (Varley et al., 2024).

Nevertheless, we should note that interpretation of our scalp-level spatial patterns requires caution. First, despite excellent goodness of fit in all clusters, we observed small but significant cluster differences in fit quality (see Sup. Tables 13, 14). These differences were not consistent across timepoints and did not systematically align with regional findings, making it unlikely that they fully account for the patterns reported. Moreover, the immature skull properties at this age, particularly related to the fontanel, may influence amplitude-dependent measures (Flemming et al., 2005), such as aperiodic offset. Future studies incorporating realistic head models and source-level EEG analyses may help to better account for these effects and refine regional interpretations. The anatomical basis of this apparent functional precedence of frontal regions remains to be clarified, although the observed pronounced maturational changes in posterior regions may reflect established anatomical gradients during infancy.

Actually, a more rapid maturation of primary sensory and motor regions is generally described centrally and posteriorly, in comparison with association regions more anteriorly (Dubois et al., 2015; Ouyang et al., 2019). Consistent with this early regional heterogeneity, studies of later infancy have reported larger posterior aperiodic offset (from 2 months to 4 years of age), an earlier decline in posterior theta power after 9-12 months of age, alongside less-regionally-uniform trajectories for aperiodic exponent (Rico-Pico et al., 2023; Wilkinson et al., 2024). Collectively, these findings suggest that aperiodic and oscillatory activity follow different developmental trajectories: aperiodic activity already displays regional heterogeneity between TEA and 2mCA, whereas periodic theta power seems more spatially uniform at this stage, probably differentiating only later in infancy. These observations highlight the value of regional analyses for understanding early brain development.

### 4.3 Impact of early adversity on brain activity maturation

First, we did not detect any statistically significant differences in EEG metrics between preterm and full-term infants at any age. This was unexpected, as extreme prematurity has previously been associated with alterations in EEG spectral power and functional connectivity (Guyer et al., 2019; Tokariev et al., 2019; Yrjölä et al., 2022). In the present study, we did not observe such differences in either aperiodic or periodic metrics. The absence of a preterm vs full-term group difference requires replication, particularly in light of the limited full-term sample (n = 13 vs n = 40 preterm). It should also be noted that this sample size did not permit a fully data-driven analysis (e.g., exploring effects beyond the predefined anterior/central/posterior clusters). As a result, spatially localized effects that do not conform to this hypothesis-driven regional analysis may have been overlooked, as suggested by the topographical distributions (see Figure 3). It should nevertheless be noted that the full-term cohort showed more uniform clinical characteristics than the preterm cohort (e.g. narrower range of gestational ages). This relative homogeneity may partially compensate for the smaller full-term sample size, although replication in larger and equally well-characterized full-term cohorts will be needed.

Notably, our previous work in this same cohort identified differences in brain activity dynamics between preterm and full-term infants (Adibpour et al., 2025). While this appears to contrast with the present findings, it may reflect differences in the sensitivity of the metrics used to capture early neural alterations, arising from distinct underlying mechanisms. It is also relevant to consider that while prematurity did not impact our spectral measures at local-level, it might still disrupt network-level connectivity and dynamics (Adibpour et al., 2025; Tokariev et al., 2019; Yrjölä et al., 2022). Given diffuse white matter alterations commonly reported in preterm born infants (Kline et al., 2021; Volpe, 2009), it is possible that prematurity disrupts how distributed regions coordinate their rhythmic activity (ie. inter-regional synchronization patterns), even when each region individually generates comparable local amount of aperiodic or periodic activity in preterm and fullterm infants. Thus, our local spectral parameterization metrics may not sufficiently capture broader network-level alterations reflected for example through inter-regional connectivity measures. Extending the parameterization frameworks to the network level analyses of brain activity may therefore help delineate inter-regional synchrony driven by genuine oscillatory activity, once the variability in oscillatory frequencies is identified across infants.

Importantly, between-group comparisons might also have been influenced by heterogeneity within the preterm population. To address this, we examined variability in EEG metrics at TEA in relation to key biological and perinatal risk factors. These analyses were exploratory, given the modest size of our preterm cohort (n=40 at TEA) and the resulting subgroup imbalances detailed below; the findings should therefore be interpreted as preliminary and will require confirmation in larger samples. We first observed a sex effect on theta power (n=21 males vs n=18 females), with males showing lower theta power than females. As theta power increased with maturation in our cohort, this finding suggests lower or delayed maturation in males, consistent with our prior observations of slower EEG dynamics (Adibpour et al., 2025) and broader evidence of increased male vulnerability and poorer outcomes in infants born preterm (Hintz et al., 2006)

Second, infants born at lower gestational ages (<28 weeks) exhibited lower aperiodic exponents than those born later in our cohort (n=13, 17 and 10 across the 24–26, 26–28 and 28–32 weeks subgroups, respectively). Given that the exponent increased with maturation, this suggests that a delayed maturation associated with very early birth may persist until TEA. Of note, this contrasted with prior work in lower-risk preterm infants, largely born after 32 weeks, which reported no impact of gestational age at birth on exponent at TEA (Shuffrey et al., 2022).

We also found higher theta power at TEA in infants with small weight at birth for gestational age at birth. This result appeared counterintuitive and should therefore be interpreted cautiously. In our cohort, only 7 of the 39 infants with an identifiable theta peak had small weight for gestational age at birth, which limited statistical power for testing potential interactions between weight risk and other risk factors. This subgroup imbalance was more pronounced than for the rest of the clinical factors examined here. Nonetheless, qualitative inspection of these few infants indicated no specific effects of other clinical factors. While altered spectral power has been reported in growth-restricted neonates (Castro Conde et al., 2020; Cohen et al., 2018; Stevenson et al., 2022), larger samples are needed to determine whether our findings might be biased or reflect compensatory or atypical developmental mechanisms.

Finally, infants with a history of invasive mechanical ventilation (n=16 with ventilation vs n=24 without ventilation) showed reduced aperiodic exponent and offset at TEA, suggesting a delayed maturation in brain functioning. The direction of this effect interpretable as relatively delayed background-activity maturation, aligns well with the literature that has linked prolonged ventilation and bronchopulmonary dysplasia with immature EEG background activity, reduced spectral power (Hahn & Tharp, 1990), decreased cortical volumes and compromised white matter integrity (Elbaz et al., 2026; Kidokoro et al., 2013). Altogether, this highlights that infants with such respiratory complications are particularly sick and at risk of impaired development of early brain activity.

In conclusion, this study demonstrates that EEG spectral parameterization combined with spatial analysis provides a sensitive framework for capturing the early postnatal maturation of brain activity in preterm and full-term infants. By identifying preliminary associations with clinical risk factors and cortical microstructure, our findings suggest that separating aperiodic and periodic activity yields novel insights into functional brain development and alterations, with specific neural vulnerabilities associated with perinatal adversity.

## Acknowledgments

A.G.C. was supported by a DIM C-BRAINS PhD fellowship (2023–2026), L.D. by an ED3C PhD fellowship (2020–2024), and L.G. by an ED3CH PhD fellowship (2023–2026). S.N. received a postdoctoral fellowship from the Bettencourt Schueller Foundation, and P.A. was supported by the UK Government Horizon Europe funding guarantee scheme of MSCA (grant EP/X021947/1).This work was further supported by the French National Agency for Research (ANR grant ANR-20-CE17-0014-03 PremaLocom), the Fondation Médisite (under the aegis of the Fondation de France, grant FdF-18-00092867), the IdEx Université de Paris (ANR-18-IDEX-0001) and the French government as part of the France 2030 program (grant ANR-23-IAHU-0010, IHU Robert-Debré du Cerveau de l’Enfant).

The authors would like to thank Guillaume Dumas and the SoNeTAA platform team (funded by Fondation de France), as well as clinical teams at Robert-Debré Hospital (in particular Nathalie Medrano, Séverine Baron, Mireille Toquer and Magali Riche) and at NeuroSpin/UNIACT (in particular Gaëlle Mediouni and Bernadette Martins) whose collaboration made it possible to conduct this study in a clinical setting. We are also grateful for helpful discussions with Catherine Chiron, Anna Kaminska, Claire Kabdebon, Fabrice Wallois and Nadège Roche-Labarbe. Last but not least, we would like to sincerely thank all the infants and parents who participated in this study.

---

**Supplementary Table 1.** Comparison of demographic and clinical characteristics between preterm infants with recordings at Session 1 (TEA) only (attrition) and those with longitudinal data at both Sessions (TEA + 2 mCA)

| Variable | Session 1 only (N = 25) | Session 1 and 2 (N = 15) | p-value |
| --- | --- | --- | --- |
| **Sex** |  |  |  |
| Male | 14 (56.0%) | 8 (53.3%) | 1.00 |
| Female | 11 (44.0%) | 7 (46.7%) |  |
| **Low birth weight risk** |  |  |  |
| Yes | 5 (20.0%) | 2 (13.3%) | 0.69 |
| No | 20 (80.0%) | 13 (86.7%) |  |
| **Invasive ventilation** |  |  |  |
| Yes | 10 (40.0%) | 6 (40.0%) | 1.00 |
| No | 15 (60.0%) | 9 (60.0%) |  |
| **Age group at EEG** |  |  |  |
| G1 | 9 (36.0%) | 4 (26.7%) | 0.60 |
| G2 | 9 (36.0%) | 8 (53.3%) |  |
| G3 | 7 (28.0%) | 3 (20.0%) |  |
| **Age at EEG (weeks)** |  |  |  |
| Mean (SD) [Range] | 40.89 (0.90) [38.40 - 42.00] | 41.03 (0.77) [39.10 - 42.10] | 0.65 |

Categorical clinical variables are reported as n (%) and compared using Fisher's exact tests. PMA at EEG is reported as mean (SD) and range and compared using Mann-Whitney. No significant differences were observed between groups on any variable (all $p \geq 0.60$ ).

**Supplementary Table 2.** Goodness of fit (R²) and Error from Models #1 and #2 for global power spectrum.

| Group | Session | Model | $R^2$ (mean ± SD) | Absolute error (mean ± SD) |
| --- | --- | --- | --- | --- |
| Preterms | TEA | Model #1 | $0.997 \pm 0.003$ | $0.030 \pm 0.012$ |
| | | Model #2 | $0.998 \pm 0.002$ | $0.023 \pm 0.007$ |
| | 2mCA | Model #1 | $0.993 \pm 0.005$ | $0.055 \pm 0.018$ |
| | | Model #2 | $0.996 \pm 0.002$ | $0.032 \pm 0.008$ |
| Fullterms | 0m | Model #1 | $0.997 \pm 0.003$ | $0.032 \pm 0.015$ |
| | | Model #2 | $0.998 \pm 0.001$ | $0.022 \pm 0.003$ |
| | 2m | Model #1 | $0.992 \pm 0.002$ | $0.062 \pm 0.010$ |
| | | Model #2 | $0.995 \pm 0.002$ | $0.035 \pm 0.009$ |

*Model #1 was fitted on the 0.5-30 Hz range and Model #2 was fitted on the 0.5-10 Hz range. Abbreviations: TEA = term-equivalent age; 0/2m(CA) = 0/2 months (corrected age).*

**Supplementary Table 3.** Theta peak center frequency from Model #1 (0.5- 30 Hz) and Model #2 (0.5-10 Hz), across age groups and clusters.

| Group | Session | Region | N | Mean center frequency Model #1 | Mean center frequency Model #2 | Statistical Test | p-value |
| --- | --- | --- | --- | --- | --- | --- | --- |
| Preterms | TEA | Global | 30 | 4,79 | 4,79 | Wilcoxon | 0,61 |
|  |  | Anterior | 31 | 4,68 | 4,70 | Wilcoxon | 0,25 |
|  |  | Central | 27 | 4,94 | 4,97 | Wilcoxon | 0,93 |
|  |  | Posterior | 28 | 4,75 | 4,78 | Wilcoxon | 0,08 |
|  | 2mCA | Global | 14 | 4,54 | 4,51 | Wilcoxon | 1,00 |
|  |  | Anterior | 10 | 4,70 | 4,52 | Wilcoxon | 1,00 |
|  |  | Central | 11 | 4,66 | 4,68 | Wilcoxon | 1,00 |
|  |  | Posterior | 13 | 4,56 | 4,64 | Wilcoxon | 0,50 |
| Fullterms | 0mCA | Global | 9 | 4,67 | 4,82 | Wilcoxon | 0,73 |
|  |  | Anterior | 9 | 4,87 | 4,83 | Wilcoxon | 0,65 |
|  |  | Central | 9 | 4,84 | 4,87 | Paired t-test | 0,65 |
|  |  | Posterior | 6 | 4,79 | 4,71 | Paired t-test | 0,32 |
|  | 2mCA | Global | 8 | 4,70 | 4,84 | Wilcoxon | 0,95 |
|  |  | Anterior | 7 | 4,63 | 4,74 | Paired t-test | 0,11 |
|  |  | Central | 7 | 4,86 | 4,54 | Wilcoxon | 0,22 |
|  |  | Posterior | 6 | 5,05 | 4,88 | Paired t-test | 0,12 |

*Model #1 was fitted on the 0.5-30 Hz range and Model #2 was fitted on the 0.5-10 Hz range.*
*Abbreviations: TEA = term-equivalent age; 0/2m(CA) = 0/2 months (corrected age).*
*N = number of infants with a valid theta peak in both models for a given region.*
*The choice between paired t-test and Wilcoxon signed-rank test was determined by the Shapiro–Wilk normality checks.*

**Supplementary Table 4.**
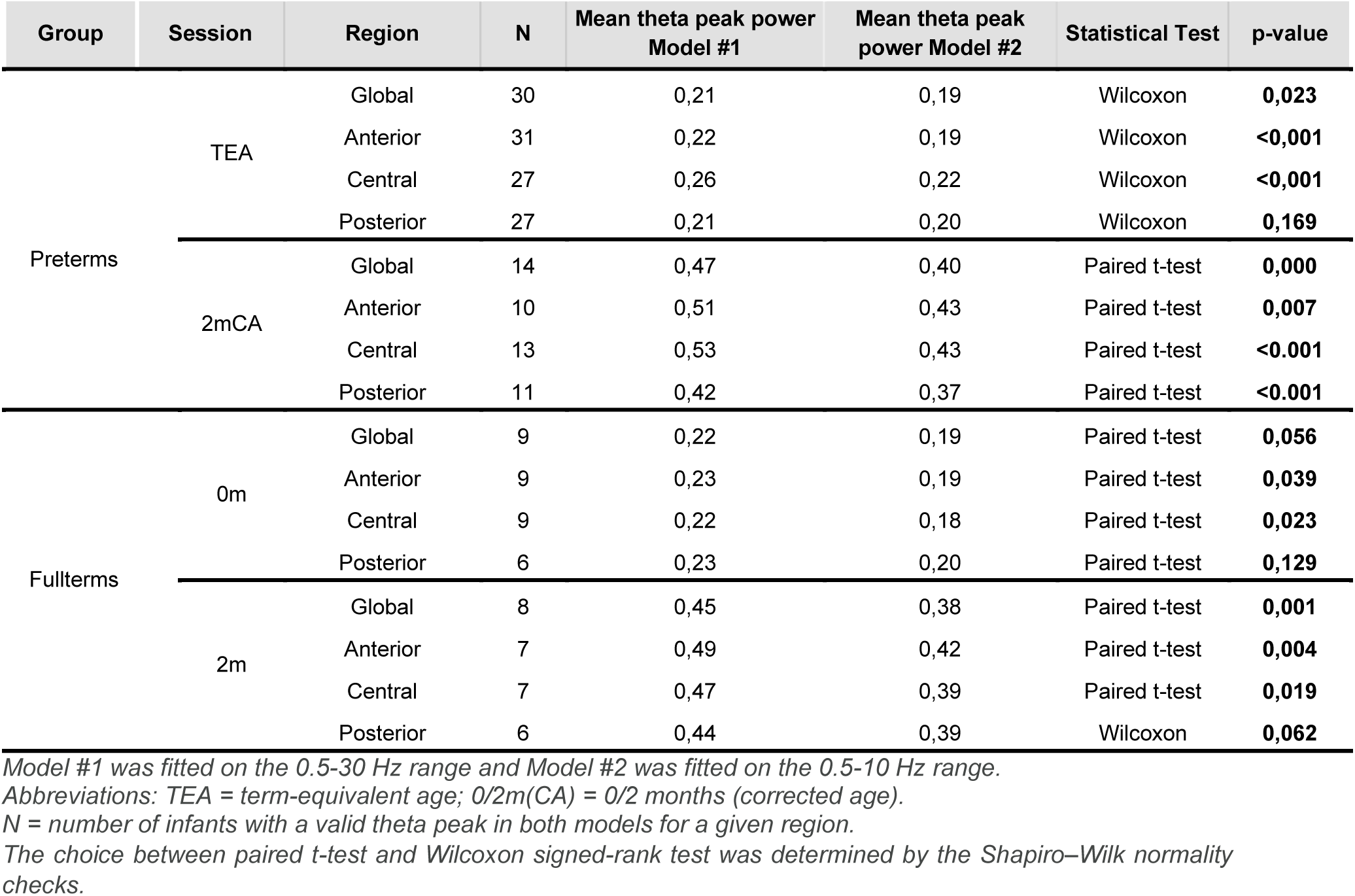
Paired comparisons of theta peak power estimates between Model #1 and Model #2, across age groups and clusters.

**Supplementary Table 5.**
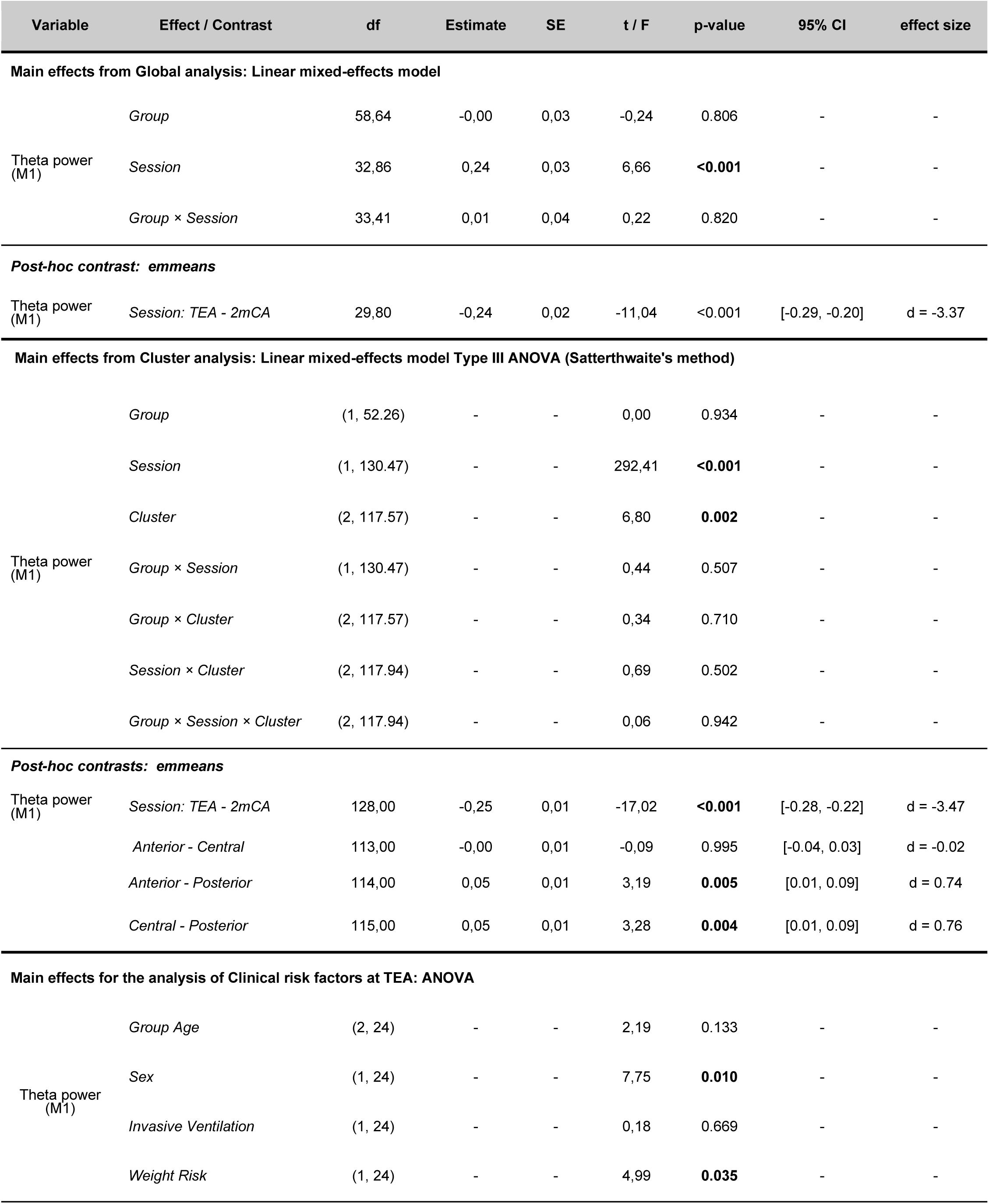
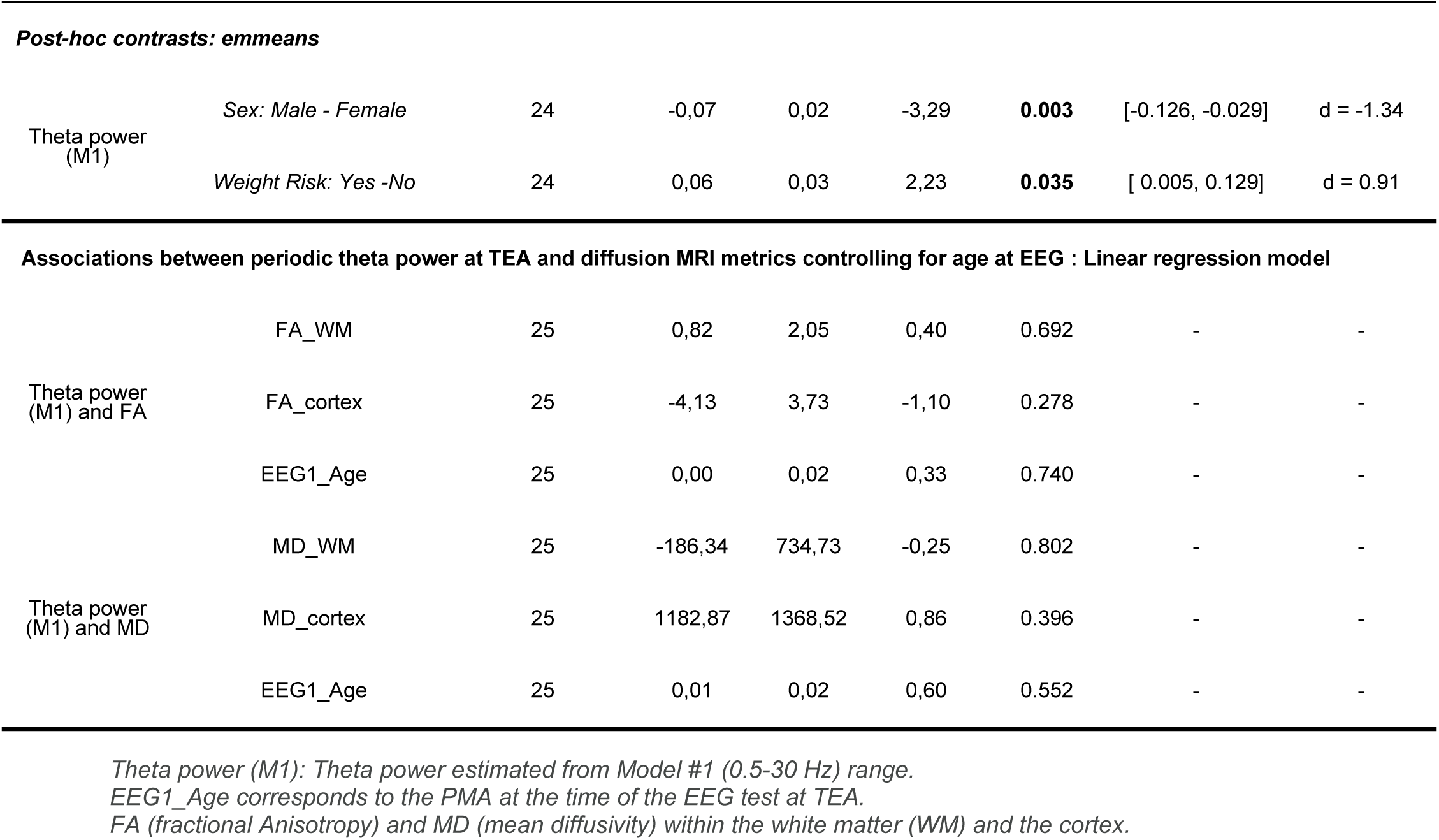
Summary of main effects and posthoc tests for theta power estimates obtained from Model #1 (0.5-30 Hz) for subjects with a valid theta peak in the broadband fit (0.5-30 Hz)

| Variable | Effect / Contrast | df | Estimate | SE | t / F | p-value | 95% CI | effect size |
| --- | --- | --- | --- | --- | --- | --- | --- | --- |
| **Main effects from Global analysis: Linear mixed-effects model** |  |  |  |  |  |  |  |  |
| Theta power (M1) | Group | 58,64 | -0,00 | 0,03 | -0,24 | 0.806 | - | - |
|  | Session | 32,86 | 0,24 | 0,03 | 6,66 | **<0.001** | - | - |
|  | Group × Session | 33,41 | 0,01 | 0,04 | 0,22 | 0.820 | - | - |
| **Post-hoc contrast: emmeans** |  |  |  |  |  |  |  |  |
| Theta power (M1) | Session: TEA - 2mCA | 29,80 | -0,24 | 0,02 | -11,04 | <0.001 | [-0.29, -0.20] | d = -3.37 |
| **Main effects from Cluster analysis: Linear mixed-effects model Type III ANOVA (Satterthwaite's method)** |  |  |  |  |  |  |  |  |
| Theta power (M1) | Group | (1, 52.26) | - | - | 0,00 | 0.934 | - | - |
|  | Session | (1, 130.47) | - | - | 292,41 | **<0.001** | - | - |
|  | Cluster | (2, 117.57) | - | - | 6,80 | **0.002** | - | - |
|  | Group × Session | (1, 130.47) | - | - | 0,44 | 0.507 | - | - |
|  | Group × Cluster | (2, 117.57) | - | - | 0,34 | 0.710 | - | - |
|  | Session × Cluster | (2, 117.94) | - | - | 0,69 | 0.502 | - | - |
|  | Group × Session × Cluster | (2, 117.94) | - | - | 0,06 | 0.942 | - | - |
| **Post-hoc contrasts: emmeans** |  |  |  |  |  |  |  |  |
| Theta power (M1) | Session: TEA - 2mCA | 128,00 | -0,25 | 0,01 | -17,02 | **<0.001** | [-0.28, -0.22] | d = -3.47 |
|  | Anterior - Central | 113,00 | -0,00 | 0,01 | -0,09 | 0.995 | [-0.04, 0.03] | d = -0.02 |
|  | Anterior - Posterior | 114,00 | 0,05 | 0,01 | 3,19 | **0.005** | [0.01, 0.09] | d = 0.74 |
|  | Central - Posterior | 115,00 | 0,05 | 0,01 | 3,28 | **0.004** | [0.01, 0.09] | d = 0.76 |
| **Main effects for the analysis of Clinical risk factors at TEA: ANOVA** |  |  |  |  |  |  |  |  |
| Theta power (M1) | Group Age | (2, 24) | - | - | 2,19 | 0.133 | - | - |
|  | Sex | (1, 24) | - | - | 7,75 | **0.010** | - | - |
|  | Invasive Ventilation | (1, 24) | - | - | 0,18 | 0.669 | - | - |
|  | Weight Risk | (1, 24) | - | - | 4,99 | **0.035** | - | - |
| **Post-hoc contrasts: emmeans** |  |  |  |  |  |  |  |  |
| Theta power (M1) | Sex: Male - Female | 24 | -0,07 | 0,02 | -3,29 | **0.003** | [-0.126, -0.029] | d = -1.34 |
|  | Weight Risk: Yes -No | 24 | 0,06 | 0,03 | 2,23 | **0.035** | [ 0.005, 0.129] | d = 0.91 |
| Theta power (M1) and FA | FA_WM | 25 | 0,82 | 2,05 | 0,40 | 0.692 | - | - |
|  | FA_cortex | 25 | -4,13 | 3,73 | -1,10 | 0.278 | - | - |
|  | EEG1_Age | 25 | 0,00 | 0,02 | 0,33 | 0.740 | - | - |
| Theta power (M1) and MD | MD_WM | 25 | -186,34 | 734,73 | -0,25 | 0.802 | - | - |
|  | MD_cortex | 25 | 1182,87 | 1368,52 | 0,86 | 0.396 | - | - |
|  | EEG1_Age | 25 | 0,01 | 0,02 | 0,60 | 0.552 | - | - |

*Theta power (M1): Theta power estimated from Model #1 (0.5-30 Hz) range.*
*EEG1\_Age corresponds to the PMA at the time of the EEG test at TEA.*
*FA (fractional Anisotropy) and MD (mean diffusivity) within the white matter (WM) and the cortex.*

**Supplementary Table 6.** Summary of main effects from statistical models of global measures (cross-sectional cohort)

| Variable | Effect-Contrast | df | Estimate | SE | t-value | Pr(>\|t\|) |
| --- | --- | --- | --- | --- | --- | --- |
| Aperiodic offset | Group | 73 | -0.01 | 0.08 | -0.17 | 0.859 |
|  | Session | 73 | 0.39 | 0.10 | 3.59 | **<0.001** |
|  | Group*Session | 73 | 0.03 | 0.13 | 0.29 | 0.769 |
| Aperiodic exponent | Group | 73 | -0.01 | 0.05 | -0.18 | 0.854 |
|  | Session | 73 | 0.05 | 0.07 | 0.73 | 0.465 |
|  | Group*Session | 73 | 0.03 | 0.08 | 0.37 | 0.713 |
| Theta power | Group | 68 | 0.00 | 0.02 | 0.21 | 0.827 |
|  | Session | 34 | 0.20 | 0.02 | 6.91 | **<0.001** |
|  | Group*Session | 35 | 0.01 | 0.03 | 0.35 | 0.722 |

*Linear mixed model results for metrics obtained from global cluster. Sample size: at TEA: n=40 in preterm group; n=11 in full-term group; at 2mCA: n=16 in preterm group and n=10 in full-term group. P-values for significant effects are highlighted in bold.*

**Supplementary Table 7.** Summary of main effects from statistical models of global measures (longitudinal cohort)

| Variable | Effect- Contrast | df | Estimate | SE | t-value | Pr(>\|t\|) |
| --- | --- | --- | --- | --- | --- | --- |
| Aperiodic offset | Group | 42 | 0.05 | 0.10 | 0.58 | 0.559 |
|  | Session | 42 | 0.47 | 0.11 | 4.08 | **<0.001** |
|  | Group*Session | 42 | -0.10 | 0.14 | -0.70 | 0.482 |
| Aperiodic exponent | Group | 42 | 0.03 | 0.07 | 0.47 | 0.638 |
|  | Session | 42 | 0.08 | 0.08 | 1.00 | 0.321 |
|  | Group*Session | 42 | -0.04 | 0.10 | -0.40 | 0.689 |
| Theta power | Group | 31 | 0.05 | 0.03 | 1.56 | 0.128 |
|  | Session | 18 | 0.22 | 0.03 | 6.49 | **<0.001** |
|  | Group*Session | 18 | -0.03 | 0.04 | -0.81 | 0.424 |

*Linear mixed effect results reported from ANOVA tables for spectral metrics obtained from global cluster. Longitudinal sample size n=15 preterms; n=8 full-terms. P-values for significant effects are highlighted in bold.*

**Supplementary Table 8.** Summary of post-hoc tests for the significant effects of models on global measures (longitudinal cohort)

| Variable | Effect- Contrast | df | Estimate | SE | t ratio | p- value | confidence interval | effect size |
| --- | --- | --- | --- | --- | --- | --- | --- | --- |
| Offset | Effect of Session: |  |  |  |  |  |  |  |
|  | TEA - 2mCA | 21 | -0.42 | 0.07 | -5.88 | **<0.001** | [-0.57, -0.27] | -1.82 |
| Theta power | Effect of Session: |  |  |  |  |  |  |  |
|  | TEA - 2mCA | 18 | -0.20 | 0.02 | -9.65 | **<0.001** | [-0.25, -0.16] | -3.20 |

*Sample size in longitudinal model: n= 15 in preterm group and n= 8 in full-term group. P-values for significant effects are highlighted in bold.*

**Supplementary Table 9.** Summary of main effects from statistical models of cluster-based measures (cross-sectional cohort)

| Variable | Effect- Contrast | df | F-value | Pr(>F) |
| --- | --- | --- | --- | --- |
| Aperiodic offset | Group | 49.55 | 0.16 | 0.684 |
|  | Session | 183.58 | 211.31 | **<0.001** |
|  | Cluster | 164.52 | 8.37 | **<0.001** |
|  | Group*Session | 183.58 | 1.73 | 0.188 |
|  | Group*Cluster | 164.52 | 1.10 | 0.334 |
|  | Session*Cluster | 164.52 | 8.29 | **<0.001** |
|  | Group*Session*Cluster | 164.52 | 0.31 | 0.732 |
| Aperiodic exponent | Group | 49.23 | 0.21 | 0.642 |
|  | Session | 192.27 | 10.63 | **0.001** |
|  | Cluster | 164.76 | 0.37 | 0.685 |
|  | Group*Session | 192.27 | 0.55 | 0.457 |
|  | Group*Cluster | 164.79 | 0.78 | 0.459 |
|  | Session*Cluster | 164.79 | 6.49 | **0.001** |
|  | Group*Session*Cluster | 164.79 | 0.27 | 0.757 |
| Theta power | Group | 52.16 | 0.10 | 0.744 |
|  | Session | 178.38 | 445.57 | **<0.001** |
|  | Cluster | 157.93 | 0.96 | 0.381 |
|  | Group*Session | 178.38 | 1.60 | 0.206 |
|  | Group*Cluster | 157.93 | 1.25 | 0.287 |
|  | Session*Cluster | 157.93 | 0.21 | 0.803 |
|  | Group*Session*Cluster | 157.93 | 0.28 | 0.754 |

Linear mixed model results reported from ANOVA tables for cluster-based measures. Sample size: at TEA: n=40 in preterm group; n=11 in full-term group at TEA; at 2mCA: n=16 in preterm group and n=10 in full-term group. P-values for significant effects are highlighted in bold

**Supplementary Table 10.** Summary of main effects from statistical models of cluster-based measures (longitudinal cohort)

| Variable | Effect- Contrast | df | F-value | Pr(>F) |
| --- | --- | --- | --- | --- |
| Aperiodic offset | Group | 21 | 0.00 | 0.995 |
|  | Session | 105 | 140.65 | **<0.001** |
|  | Cluster | 105 | 4.21 | **0.017** |
|  | Group*Session | 105 | 2.04 | 0.155 |
|  | Group*Cluster | 105 | 1.51 | 0.225 |
|  | Session*Cluster | 105 | 3.45 | **0.035** |
|  | Group*Session*Cluster | 105 | 0.23 | 0.793 |
| Aperiodic exponent | Group | 21 | 0.11 | 0.734 |
|  | Session | 105 | 5.87 | **0.017** |
|  | Cluster | 105 | 0.20 | 0.818 |
|  | Group*Session | 105 | 0.82 | 0.366 |
|  | Group*Cluster | 105 | 1.11 | 0.332 |
|  | Session*Cluster | 105 | 2.40 | 0.095 |
|  | Group*Session*Cluster | 105 | 0.33 | 0.715 |
| Theta power | Group | 20.33 | 0.57 | 0.456 |
|  | Session | 90.38 | 298.22 | **<0.001** |
|  | Cluster | 92.13 | 0.180 | 0.835 |
|  | Group*Session | 90.38 | 2.13 | 0.147 |
|  | Group*Cluster | 92.13 | 0.72 | 0.486 |
|  | Session*Cluster | 90.38 | 0.45 | 0.634 |
|  | Group*Session*Cluster | 90.38 | 0.26 | 0.770 |

*Linear mixed model results reported from ANOVA tables for cluster-based measures. Sample size in longitudinal model n=15 in preterm group; n=8 in full-term group. P-values for significant effects are highlighted in bold.*

**Supplementary Table 11.** Summary of post-hoc tests for significant effects of models on cluster measures (longitudinal data)

| Variable | Effect- Contrast | df | Estimate | SE | t ratio | p- value | confidence interval | effect size |
| --- | --- | --- | --- | --- | --- | --- | --- | --- |
| **Offset** | *Effect of Session:* |  |  |  |  |  |  |  |
| TEA - 2mCA | All clusters | 105 | -0.44 | 0.03 | -11.86 | **<0.001** | [-0.51, -0.36] | -2.12 |
|  | *Effect of Cluster:* |  |  |  |  |  |  |  |
| TEA and 2mCA | Anterior - Central | 105 | 0.13 | 0.04 | 2.90 | **0.01** | [0.02, 0.24] | 0.63 |
| TEA and 2mCA | Anterior - Posterior | 105 | 0.06 | 0.04 | 1.44 | 0.32 | [-0.04, 0.17] | 0.31 |
| TEA and 2mCA | Central - Posterior | 105 | -0.06 | 0.04 | -1.46 | 0.31 | [-0.17, 0.04] | -0.32 |
|  | *Effect of Session\*Cluster:* |  |  |  |  |  |  |  |
| TEA - 2mCA | Anterior Cluster | 105 | -0.31 | 0.06 | -4.86 | **<0.001** | [-0.44, -0.18] | -1.51 |
| TEA - 2mCA | Central Cluster | 105 | -0.46 | 0.06 | -7.13 | **<0.001** | [-0.59, -0.33] | -2.21 |
| TEA - 2mCA | Posterior Cluster | 105 | -0.55 | 0.06 | -8.54 | **<0.001** | [-0.68, -0.42] | -2.65 |
| TEA | Anterior - Central | 105 | 0.20 | 0.06 | 3.18 | **0.005** | [0.05, 0.36] | 0.98 |
| TEA | Anterior - Posterior | 105 | 0.18 | 0.06 | 2.86 | **0.013** | [0.03, 0.33] | 0.88 |
| TEA | Central - Posterior | 105 | -0.02 | 0.06 | -0.32 | 0.943 | [-0.17, 0.13] | -0.10 |
| 2mCA | Anterior - Central | 105 | 0.05 | 0.06 | 0.91 | 0.629 | [-0.09, 0.21] | 0.28 |
| 2mCA | Anterior - Posterior | 105 | -0.05 | 0.06 | -0.82 | 0.690 | [-0.20, 0.10] | -0.25 |
| 2mCA | Central - Posterior | 105 | -0.11 | 0.06 | -1.74 | 0.195 | [-0.26, 0.04] | -0.53 |
| **Exponent** | *Effect of Session:* |  |  |  |  |  |  |  |
| TEA - 2mCA | All clusters | 105 | -0.06 | 0.02 | -2.42 | **0.017** | [-0.12, -0.01] | -0.43 |
| **Theta power** | *Effect of Session:* |  |  |  |  |  |  |  |
| TEA - 2mCA | All clusters | 91 | -0.21 | 0.01 | -17.26 | **<0.001** | [-0.23, -0.18] | -3.26 |

Sample size in longitudinal model: n= 15 in preterm group and n= 8 in full-term group. P-values for significant effects are highlighted in bold

**Supplementary Table 12.** Summary of main effects from statistical models of clinical risk factors in preterm infants at TEA.

| Variable | Effect- Contrast | df | F-value | Pr(>F) |
| --- | --- | --- | --- | --- |
| Aperiodic offset | Sex | 34 | 0.04 | 0.836 |
|  | Group of GA at birth | 34 | 1.00 | 0.375 |
|  | Small weight for GA at birth | 34 | 3.18 | 0.083 |
|  | Invasive ventilation | 34 | 6.95 | **0.012** |
| Aperiodic exponent | Sex | 34 | 0.54 | 0.466 |
|  | Group of GA at birth | 34 | 5.39 | **0.009** |
|  | Small weight for GA at birth | 34 | 0.02 | 0.871 |
|  | Invasive ventilation | 34 | 6.38 | **0.016** |
| Theta power | Sex | 33 | 4.68 | **0.037** |
|  | Group of GA at birth | 33 | 1.10 | 0.342 |
|  | Small weight for GA at birth | 33 | 5.34 | **0.027** |
|  | Invasive ventilation | 33 | 2.49 | 0.123 |

Sample size of preterm group at TEA: $n = 40$ . $P$ -values for significant effects are highlighted in bold.

**Supplementary Table 13.** Goodness-of-fit metrics for the spectral parameterization across sessions and clusters.

| Group | Session | n (subjects) | Cluster | R<sup>2</sup> (mean ± SD) | Fit error (mean ± SD) |
| --- | --- | --- | --- | --- | --- |
| Preterms | TEA | 40 | Anterior | 0.997 ± 0.002 | 0.028 ± 0.011 |
|  |  |  | Central | 0.997 ± 0.002 | 0.027 ± 0.009 |
|  |  |  | Posterior | 0.995 ± 0.004 | 0.035 ± 0.012 |
|  | 2mCA | 16 | Anterior | 0.989 ± 0.008 | 0.066 ± 0.025 |
|  |  |  | Central | 0.994 ± 0.005 | 0.045 ± 0.018 |
|  |  |  | Posterior | 0.994 ± 0.003 | 0.050 ± 0.018 |
| Fullterms | 0m | 11 | Anterior | 0.997 ± 0.001 | 0.027 ± 0.010 |
|  |  |  | Central | 0.996 ± 0.002 | 0.031 ± 0.012 |
|  |  |  | Posterior | 0.995 ± 0.002 | 0.037 ± 0.011 |
|  | 2m | 10 | Anterior | 0.993 ± 0.004 | 0.051 ± 0.017 |
|  |  |  | Central | 0.993 ± 0.003 | 0.053 ± 0.010 |
|  |  |  | Posterior | 0.992 ± 0.002 | 0.058 ± 0.014 |

**Supplementary Table 14.** Pairwise post-hoc regional comparisons of fit quality.

| Group | Session | Fitting parameter | Contrast | df | t | p | Cohen's d |
| --- | --- | --- | --- | --- | --- | --- | --- |
| Preterms | TEA | R<sup>2</sup> | Anterior-Central | 78 | -0.123 | 1.000 | -0.028 |
|  |  |  | Anterior- Posterior | 78 | 4.228 | **< .001** | 0.945 |
|  |  |  | Central- Posterior | 78 | 4.351 | **< .001** | 0.973 |
|  |  | Fit error | Anterior-Central | 78 | 0.583 | 1.000 | 0.130 |
|  |  |  | Anterior- Posterior | 78 | -4.172 | **< .001** | -0.933 |
|  |  |  | Central- Posterior | 78 | -4.755 | **< .0001** | -1.063 |
|  | 2mCA | R<sup>2</sup> | Anterior- Central | 30 | -2.764 | **0.029** | -0.977 |
|  |  |  | Anterior- Posterior | 30 | -2.612 | **0.041** | -0.923 |
|  |  |  | Central- Posterior | 30 | 0.152 | 1.000 | 0.054 |
|  |  | Fit error | Anterior - Central | 30 | 3.171 | **0.010** | 1.121 |
|  |  |  | Anterior- Posterior | 30 | 2.382 | 0.071 | 0.842 |
|  |  |  | Central- Posterior | 30 | -0.789 | 1.000 | -0.279 |
| Fullterms | 0m | R<sup>2</sup> | Anterior - Central | 20 | 1.634 | 0.354 | 0.697 |
|  |  |  | Anterior- Posterior | 20 | 3.441 | **0.007** | 1.467 |
|  |  |  | Central- Posterior | 20 | 1.807 | 0.257 | 0.771 |
|  |  | Fit error | Anterior - Central | 20 | -1.277 | 0.649 | -0.544 |
|  |  |  | Anterior- Posterior | 20 | -2.983 | **0.022** | -1.272 |
|  |  |  | Central- Posterior | 20 | -1.707 | 0.310 | -0.728 |
|  | 2m | R<sup>2</sup> | Anterior- Central | 18 | 0.171 | 1.000 | 0.076 |
|  |  |  | Anterior- Posterior | 18 | 0.607 | 1.000 | 0.271 |
|  |  |  | Central- Posterior | 18 | 0.436 | 1.000 | 0.195 |
|  |  | Fit error | Anterior- Central | 18 | -0.397 | 1.000 | -0.178 |
|  |  |  | Anterior- Posterior | 18 | -1.103 | 0.854 | -0.493 |
|  |  |  | Central- Posterior | 18 | -0.705 | 1.000 | -0.315 |

Pairwise comparisons performed following linear mixed-effects models with Cluster as fixed effect. Significant contrasts ( $p < .05$ ) are highlighted in bold.

## Notes

### Competing Interest Statement

The authors have declared no competing interest.

### Summary of Updates

The updated article has added a number of complementary analyses to control for some effects and changed some parts of the discussion and supplementary materials.

